# Identification and functional analysis of αKNL2 genes in cowpea

**DOI:** 10.64898/2026.07.27.740913

**Authors:** Andriy Kochevenko, Itzel Amasende-Morales, Gloria León-Martínez, Judith Lúa, Osvaldo Ruiz-Maciel, Jörg Fuchs, Jean-Philippe Vielle-Calzada, Andreas Houben

## Abstract

Although the KINETOCHORE NULL2 (KNL2) protein is an essential inner centromeri c protein that is crucially important for assembly and functioning of kinetochores, our understanding of its organization, dynamics and function of distinct isoforms in the cells of plant species undergoing mitosis/meiosis is far from complete. In this study, we identified and characterized two *αKNL2.1*/*αKNL2.2* genes in cowpea. GUS reporter constructs and qRT-PCR revealed that the expression profiles of both genes were variable across organs, with the highest expression in leaves and roots. Using an EYFP gene fusion coupled with immunostaining, it was demonstrated that both αKNL2 variants colocalized at centromeres in a cell-cycle-dependent manner. The CRISPR/Cas9 technique was used to generate various in-frame deletion and out-of-frame knock-out *αknl2* mutants. Single- and double-gene knock-out mutants were generated, and the effects of mutations on plant development and seed setting were analyzed. The results are discussed both with respect to the roles of these proteins in kinetochore assembly and in the context of using *αKNL2* genes for *in vivo* production of haploids in cowpea.

**Significance statement:** This study identifies two paralogous αKNL2 genes in cowpea and reveals their functional redundancy during centromere assembly and essential role in seed development. These findings expand our knowledge of kinetochore dynamics and provide a basis for exploring the evolutionary diversification of centromeric proteins in legumes.

## Introduction

Centromeres are specialized regions of eukaryotic chromosomes that assemble kinetochore complexes and ensure accurate chromosome segregation during cell division (Naish and Henderson 2024). The centromere-specific histone H3 variant (called CENH3 in plants and CENP-A in mammals) plays a key role in centromere specification. The presence of just a few CENP-A-containing nucleosomes is sufficient to mark the site of centromere formation (Bodor et al. 2014; Mellone and Fachinetti 2021; Naish et al. 2021; Altemose et al. 2022). CENP-A physically interacts with CENP-C and CENP-N (Allu et al. 2019; Ariyoshi et al. 2021) and recruits multiple components of the constitutive centromere-associated network (CCAN) complex of the inner kinetochore that, in turn, recruits the KMN network complex, which enables chromosomes to attach to spindle microtubules (Yatskevich et al. 2024).

Previous studies have shown that loading new CENP-A onto existing centromeres and the assembly of CENP-A nucleosomes are replication-independent (Almouzni and Cedar 2016). In fission yeast, human and plant cells, this process was reported to be restricted to either G2 (Dunleavy et al. 2007), late telophase/early G1 (Jansen et al. 2007; Dunleavy et al. 2009) or G2/prophase (Lermontova et al. 2006) phases of the cell cycle, respectively. As CENP-A loading occurs only after the next mitosis, this leads to the situation where, during centromere DNA replication, CENP-A nucleosomes are diluted to half their initial concentration in the centromeric daughter chromatin (Shelby et al. 2000; Jansen et al. 2007).

Studies in eukaryotic and yeast cells identified factors assisting CENP-A deposition at centromeres. In particular, it was reported that histone assembly factors as Holliday junction recognition protein (HJURP) in mammals, suppressor of chromosome missegregation 3 (Scm3) in yeasts and chromosome alignment defect 1 (CAL1) in flies, specifically interact with CENP-A and promote CENP-A deposition at centromeres, ensuring continuous inheritance of their position (Dunleavy et al. 2009; Foltz et al. 2009; Pidoux et al. 2009; Williams et al. 2009; Medina-Pritchard et al. 2020). HJURP binds to a CENP-A–H4 heterodimer and enables the deposition of CENP-A to the centromere in a cell cycle-dependent manner (Hu et al. 2011; Zasadzińska et al. 2013). During S-phase, HJURP can transiently associate with centromeres and, together with the minichromosome maintenance 2 (MCM2) subunit of the replicative helicase, contribute to the maintenance of epigenetic marks of centromeric chromati n during DNA replication (Huang et al. 2015; Zasadzińska et al. 2018).

In vertebrates, the recruitment of the HJURP chaperone to centromeres was reported to be controlled by the Mis18 complex, a licensing factor which specifies the site of new CENP-A nucleosome assembly (Zasadzińska et al. 2013; Pan et al. 2019). In humans, the Mis18 complex consists of Mis18a, Mis18b, and Mis18-binding protein 1 (M18BP1; also known as hsKNL2) (Fujita et al. 2007; Maddox et al. 2007). It is known that phosphorylation of the Mis18 complex mediated by cyclin-dependent kinase 1 and 2 (CDK1 and CDK2) and Polo-like kinase 1 (PLK1) controls deposition of new CENP-A (Spiller et al. 2017; Stankovic et al. 2017; Parashara et al. 2024).

In plants, the molecular mechanisms controlling the recognition and deposition of CENH3 remain poorly characterized (Takeuchi et al. 2024). Notably, the plant ortholog of a major CENP-A chaperone, HJURP, has not been identified to date (Naish and Henderson 2024).Recently, the *Arabidopsis* homolog of nuclear autoantigenic sperm protein (AtNASP) was reported to function as a CENH3 chaperone, playing a key role in *de novo* CENH3 deposition after fertilization (Le Goff et al. 2020; Takeuchi et al. 2024). Moreover, within the Mis18 complex, only the M18BP1/KNL2 protein has been found (Lermontova et al. 2013). KINETOCHORE NULL2 (KNL2) plays a key role in the deposition of new CENH3 at centromeres in plants. In angiosperms, two variants of KNL2 were described: αKNL2/βKNL2 in eudicots and γKNL2/δKNL2 in grasses (Zuo et al. 2022). These paralogous proteins differ in size, domain composition and organization, and in the presence of specific functional motifs. The N-terminus of both paralogs contains the conserved SANTA domain (∼90 amino acid residues), whereas only the αKNL2 proteins possess the CENPC-k motif at their C-terminus (Zuo et al. 2022). The N-terminal SANTA domain is not required for centromeric loading of αKNL2, whereas CENPC-k motif and two adjacent DNA-binding regions were found to be essential for successful centromeric targeting of this kinetochore protein. Moreover, it was observed that deletion of one or both DNA-binding sites substantially reduced or even prevented centromeric localization of such EYFP-tagged αKNL2-C fragments (Yalagapati et al. 2025).

*A. thaliana* αKNL2 shows centromeric localization throughout the mitotic cell cycle, except for metaphase to anaphase (Lermontova et al. 2013). A similar dynamic is observed in *Schizosaccharomyces pombe*, where Mis18 is absent from centromeres during the metaphase-anaphase transition (Hayashi et al. 2004). In humans, KNL2 is transiently recruited to centromeres following mitotic exit (Fujita et al. 2007). Knock-out of the *A. thaliana αKNL2* gene resulted in reduced levels of CENH3 protein at centromeres and 30% seed abortion in developing pods (Lermontova et al. 2013). In line with the CENH3-loading function of αKNL2, inactive *αknl2* induced the formation of haploids upon outcrossing with wild-type *A. thaliana* (Ahmadli et al. 2023). Similarly, in maize, the knock-out of KNL2 induced haploids in both male and female crosses (Li et al. 2025).

The βKNL2 protein (missing the CENPC-k motif) was also reported to be targeted to the *A. thaliana* centromere and to participate in loading CENH3. Unfortunately, the precise mechanism of its loading to the centromere is still poorly characterized. It is known that βKNL2 gene expression is considerably higher than that of its paralog, αKNL2, and, at least in some tissues, βKNL2 targeting seems to depend on αKNL2 protein (Yadala et al. 2026; Zuo et al. 2022; Yalagapati et al. 2025)

The identification of *αKNL2* as a gene harnessed to induce haploids in *A. thaliana* and maize prompted us to characterize the corresponding gene in cowpea (*Vigna unguiculata*; 2n=2×=22). Cowpea is a multipurpose legume crop mostly cultivated in the tropical and subtropical regions of Latin America, Africa, Oceania and Asia (Boukar et al. 2019). Cowpea possesses high resilience to drought and heat stress and therefore has great potential to improve agricultural sustainability and food security (Silva et al. 2024). Despite the growing importance of this crop, little is known about its centromeres. Previously, we described and functionally characterized two cowpea CENH3 homologs (Ishii et al. 2020). In this study, we provide evidence that cowpea has two αKNL2 paralogous proteins encoded by separate genes located on distinct chromosomes. Both αKNL2 isoforms were shown to be localized to centromeres. qRT-PCR and GUS reporter construct analyses demonstrated that both *αKNL2* genes are transcribed. Furthermore, CRISPR/Cas9 Mutagenesis was applied to investigate the roles of the αKNL2 genes in vegetative and reproductive development in cowpea and to test their haploidization-inducing ability.

## Results

### Cowpea encodes two variants of αKNL2

To identify the orthologous αKNL2 proteins in cowpea, BLASTP search in the Phytozome V13 database was carried out using the amino acid sequence of the known *Arabidopsis* αKNL2 protein (At5g02520) (Lermontova *et al*., 2013). As a result, three candidate *KNL2* homologous genes were identified. Deduced amino acid sequences of two of these homologs (Figure 1a; Figure S1) contained both N-terminal SANTA domain and the conserved CENPC-k motif on the C-terminal part, which are specific features of αKNL2-type proteins (Zuo et al., 2022). Therefore, all further studies were focused on these two members of the KNL2 family. These identified αKNL2 homologous protein sequences shared more than 81% sequence identity to each other and over 31% with *A. thaliana* αKNL2 (Table S1). We designated them as αKNL2.1 (Vigun11g213800) and αKNL2.2 (Vigun07g086000), respectively. Genes encoding these proteins are located on chromosomes Vu07 and Vu11, and each gene contains eight exons and seven introns (Figure 1b). Multiple sequence alignments of the predicted KNL2 proteins from cowpea and related *Fabaceae* species revealed conservation across motifs, including the N-terminal region, particularly the SANTA domain, which harbours the VxLxDW and FxxGFPxxW motifs specific to all KNL2 proteins, and the CENPC-k motif located at the C-terminal region of αKNL2 proteins (Figure S1).

**Figure 1.**
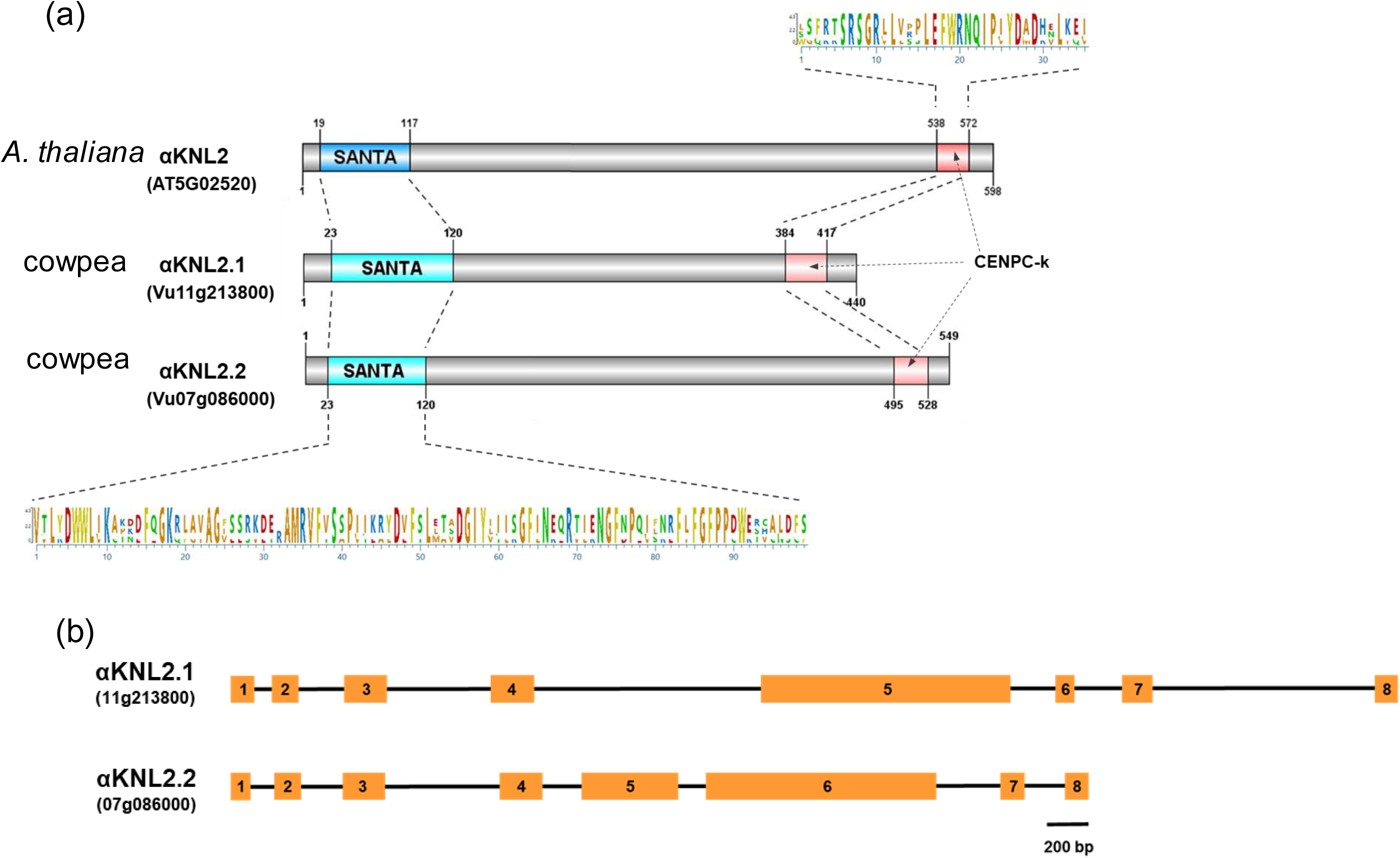
Structure of the cowpea αKNL2 genes and the encoded proteins. (a) Schematic structure of cowpea and *A. thaliana* αKNL2 proteins. The conserved protein SANTA domains and CENPC-k motifs are indicated. Amino acid positions are indicated above the sequences. (b) A schematic representation of intron/exon structure of the αKNL2 genes. The orange box represents an exon and the black solid line represents an intron.

A maximum likelihood phylogeny was generated using the inferred protein sequences of *KNL2* gene set of *Fabaceae* species. The results suggested that all predicted proteins to date could be divided into two groups: αKNL2 and βKNL2. The former group includes 14 members, while the latter includes seven (Figure S2). Moreover, the analysis indicated that the cowpea αKNL2s exhibited the closest evolutionary relationship to the corresponding protein of *Phaseolus vulgaris*.

### Expression analysis of both *αKNL2* genes

To study the expression pattern of both *αKNL2* variants in cowpea, transgenic plants bearing *αKNL2s* promoter - β-glucuronidase (GUS) gene fusion construct (proVu_αKNL2.1::GUS or proVu_αKNL2.2::GUS) were generated. Twenty independent transgenic lines for each construct expressing the GUS gene under the control of αKNL2.1 or αKNL2.2 were included in the analysis. Histochemical staining of these lines revealed that both genes were expressed in all vegetative and generative organs analyzed (Figure 2). However, substantial differences in staining intensity within the same tissues were observed for these two promoter-reporter constructs (Figure 2a1, b1; a5, b5; a9, b9). Plants bearing the proVu_αKNL2.1::GUS demonstrated stronger overall GUS expression, except for epicotyls. In this organ, no identifiable difference was detected in GUS staining intensities between the proVu_αKNL2.1::GUS and proVu_αKNL2.2::GUS lines (Figure 2a6, b6). Main differences in expression were observed in the root system of plants (Figure 2a9, b9). It was determined that *αKNL2.1* promoter provided strong expression throughout the length of the main root, except for the root tips and lateral roots. In plants expressing proVu_αKNL2.2::GUS only weak GUS activity was detected in the primary root, and no staining was observed either at root tips or lateral roots.

**Figure 2.**
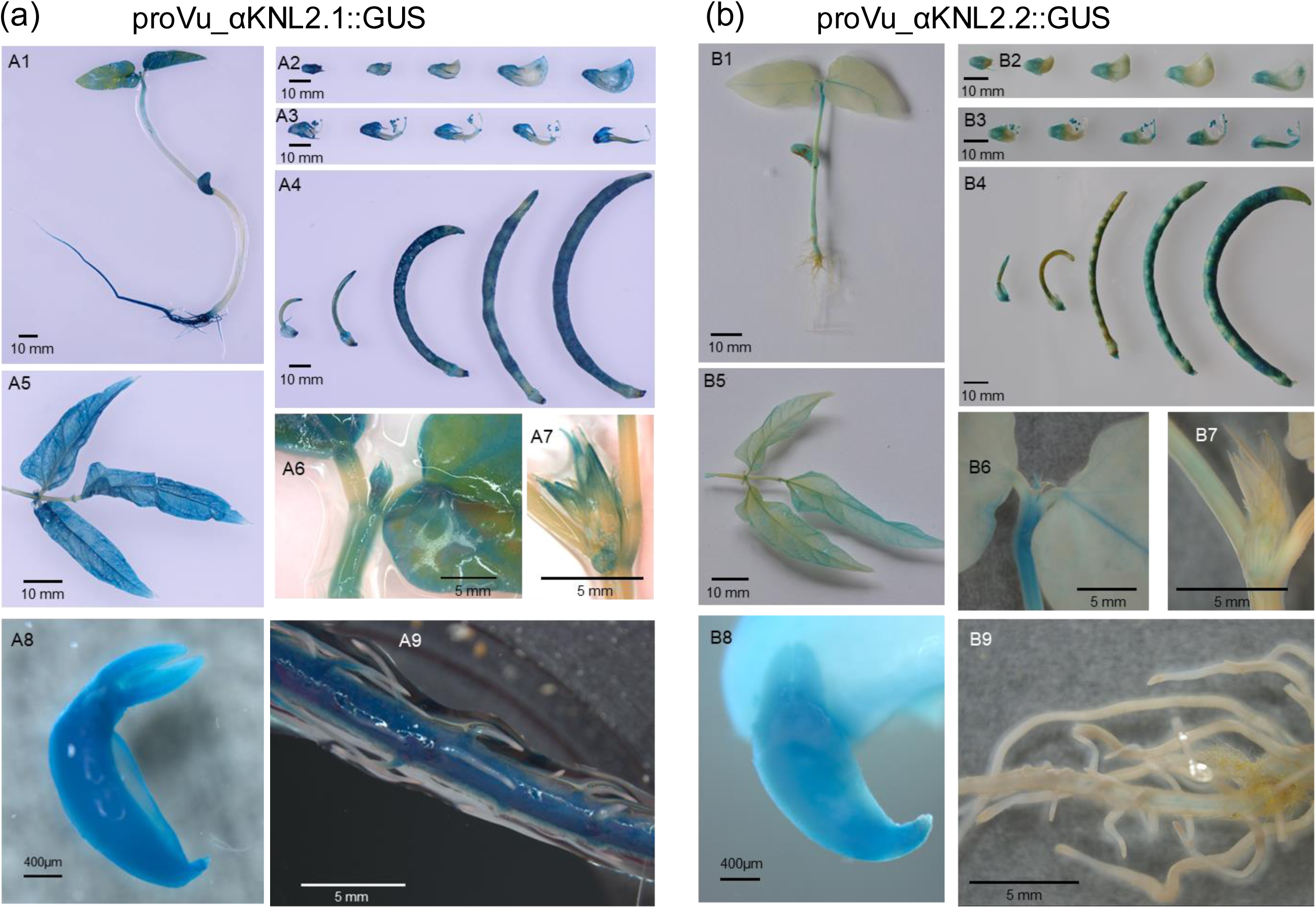
Histochemical staining of transgenic cowpea lines expressing (a) αKNL2.1:GFP-GUS and (b) αKNL2.2:GFP-GUS promoter-reporter constructs. (a) and (b), GUS activity assay of αKNL2.1 and αKNL2.2 promoters, respectively, at different development stages and organs of transgenic cowpea. Respective scale bars are represented. A1, B1 - two weeks old transgenic cowpea seedlings; A2, A3, B2, B3 - floral buds at different developmental stages; A4, B4 – pods at different developmental stages; A5, B5 – leaves; A6, B6 - epicotyls and unifoliate leaves; A7, B7 – lateral buds; A8, B8 - embryonic axis; A9, B9 – main and lateral roots.

The expression patterns of the *αKNL2* genes were examined by means of qRT-PCR analysis of RNA isolated from the roots, leaves, stems, flowers, pods and seeds. In parallel, the gene activity of cowpea CENH3.1 was determined. The results clearly demonstrated that *αKNL2.1* and *αKNL2.2* genes were expressed in all tissues analyzed. *αKNL2.1* showed constitutively high expression, while *αKNL2.2* was characterized by more variable levels across different tissues (Figure 3). Expression of *αKNL2.1* was particularly high in leaves and roots where it was comparable or even exceeded such of *CENH3.1*. The highest activity of *αKNL2.2* was detected in the leaves and flowers. At the same time, *αKNL2.2* displayed the lowest expression in the stem and pods, whereas *αKNL2.1* was, by contrast, significantly higher expressed in these tissues. In general, qRT-PCR data corroborated the previous findings with promoter-reporter constructs, indicating that αKNL2.1 had higher expression than *αKNL2.2. CENH3.1* mirrored the transcription profile of αKNL2.1.

**Figure 3.**
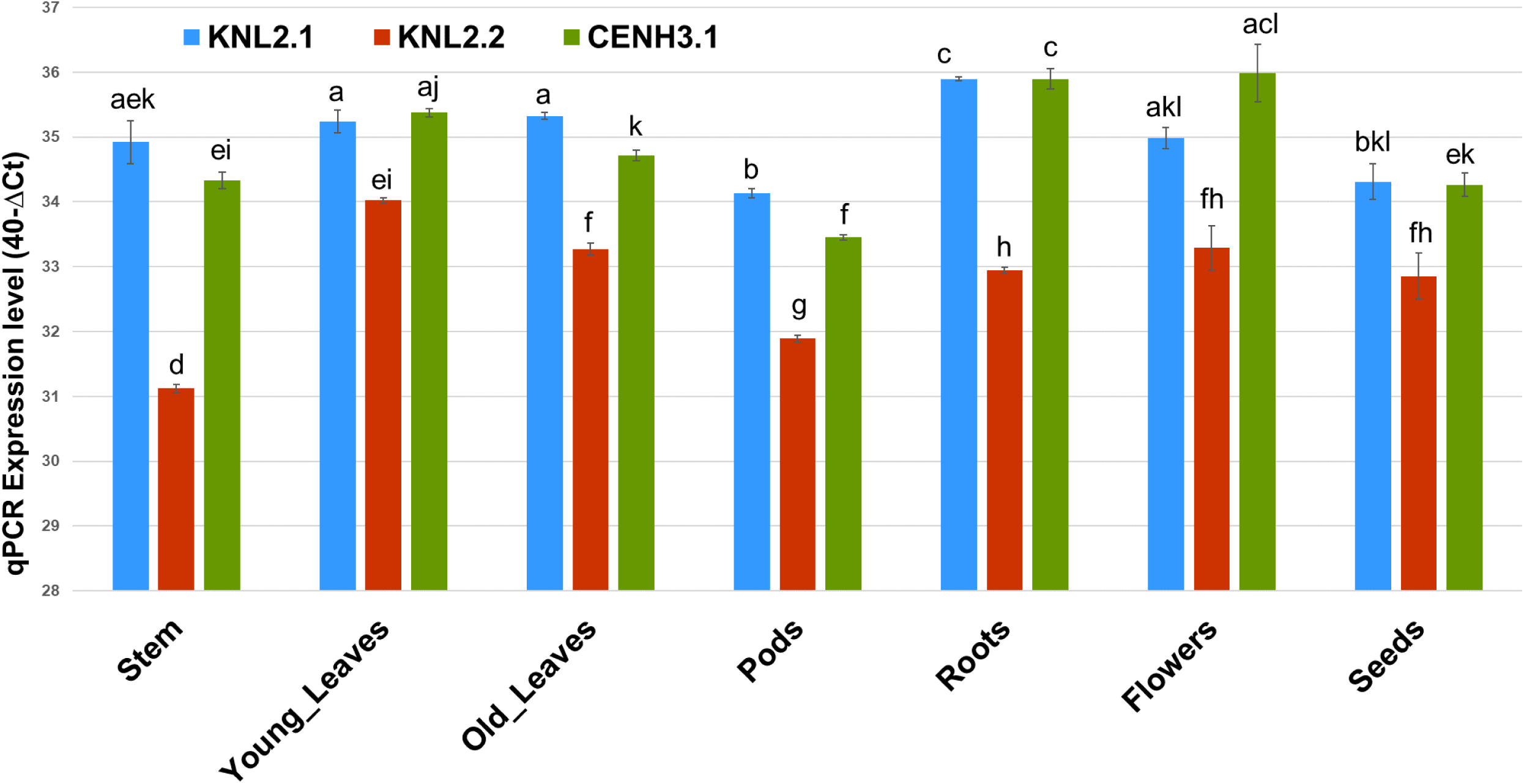
Expression profile of αKNL2 genes in vegetative and reproductive organs of cowpea. Data are means of four biological replicates and error bars represent SE. Different lowercase letters indicate significant differences (T-test, p < 0.05).

### Centromere-specific localization of both αKNL2 variants

To better understand the potential functions of αKNL2 isoforms, a subcellular localization analysis of both proteins was performed. EYFP reporter fusion constructs (EYFP_αKNL2.1_ pAGM4723 and EYFP_αKNL2.2_ pAGM4723) were transiently expressed in cowpea leaves. Observed dot-like green fluorescence signals matched with the intensely stained chromocenters of DAPI-labelled nuclei. Moreover, nuclei isolated from the tissues expressing either EYFP_αKNL2.1 or EYFP_αKNL2.2 were immunostained with anti-cowpea CENH3.1 antibodies (Figure 4a, b). Colocalization of red and green fluorescence signals corresponding to CENH3.1 and αKNL2.1/αKNL2.2, respectively, confirmed that both αKNL2 variants localized to cowpea centromeres.

**Figure 4.**
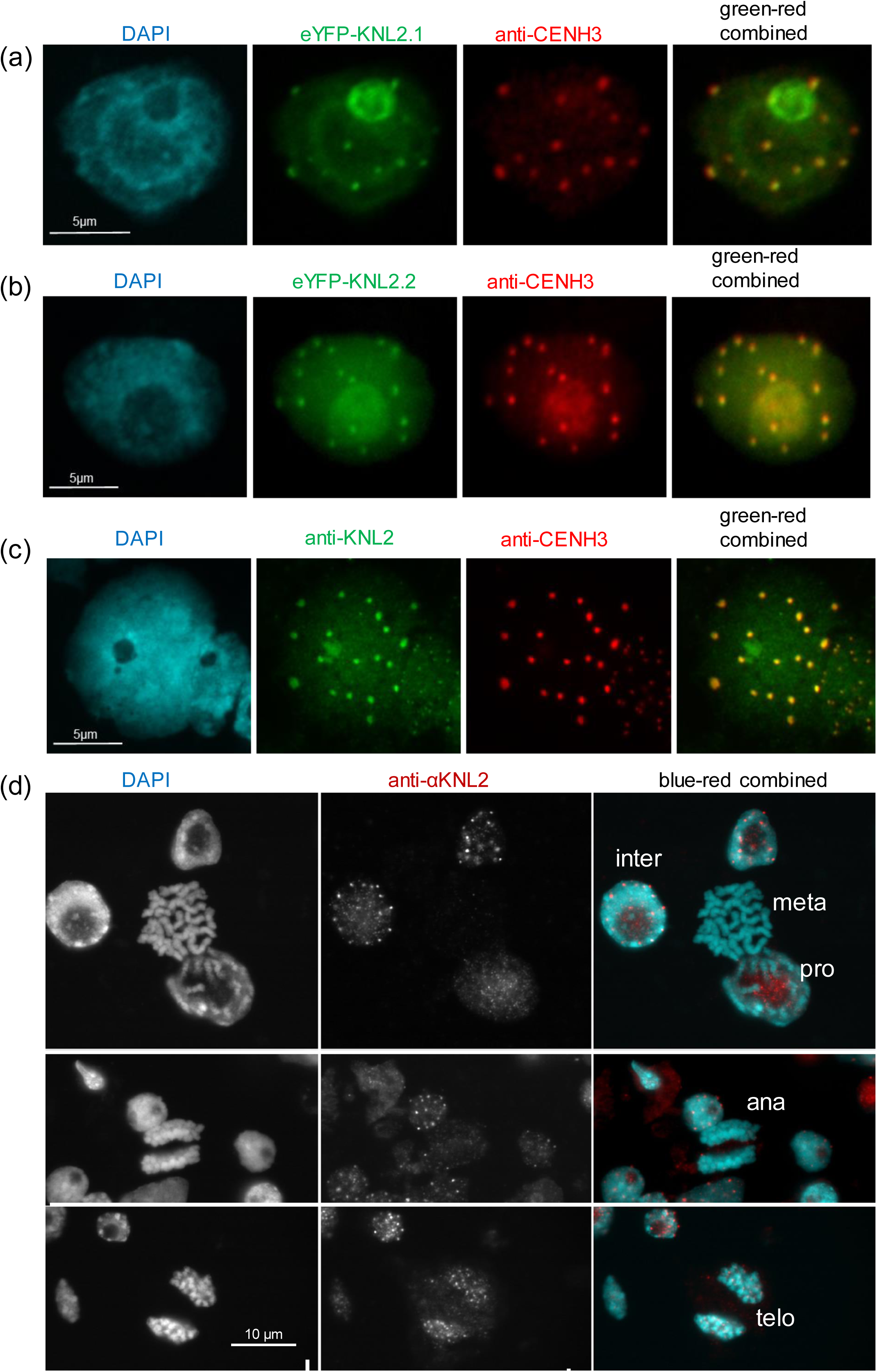
Centromeric localization of αKNL2s proteins in wild-type cowpea. (a, b) Transient expression of eYFP-αKNL2.1 and eYFP-αKNL2.2, respectively, combined with immunostaining with anti-CENH3.1 antibody. (c) Double immunostaining with anti-KNL2 and anti-CENH3.1 antibody. (d) Mitotic cells of cowpea showing anti-cowpea αKNL2 immuno signals. Note the absence of centromeric αKNL2 signals between prophase and anaphase.

To study the mitotic cell cycle dynamics of αKNL2 proteins, several rabbit polyclonal antibodies were generated against synthetic peptides corresponding to the distinct regions of both cowpea αKNL2s. However, only the antibody based on the peptide (CDVVGPKSPETPIQSQSWRQ) representing the C-terminus of αKNL2.1 produced clear dot-like immunosignals in mitotic root meristems of cowpea. The αKNL2.1 and αKNL2.2 isoforms were nearly identical in this twenty-amino-acid region, with only one amino acid substitution observed. This similarity suggests that the generated antibody does not distinguish between the αKNL2 isoforms. Double immunostaining of cowpea nuclei with anti-αKNL2 and anti-CENH3.1 antibodies confirmed the centromeric localization of αKNL2 proteins (Figure 4c). To determine the mitotic cell cycle dynamics of αKNL2, cowpea root meristematic cells were immunolabeled with anti-αKNL2. Interphase and telophase revealed centromeri c signals, while the centromeres of prophase, metaphase and anaphase chromosomes were αKNL2 negative (Figure 4d).

### Generation of CRISPR-Cas mutants

CRISPR-Cas9 technology was used to manipulate the activity of both *αKNL2* variants in cowpea. The CRISPR-P web tool (Lei et al., 2014) was used to design a set of single-guide RNAs (sgRNA) targeting the αKNL2.1/αKNL2.2 genes in cowpea (Table S2). The activity of each sgRNA was assessed using an *in vitro* cleavage assay as described by (Jinek et al. 2012) (Figure S3). Six sgRNAs demonstrating high experimental cleavage efficiencies were used to engineer Cas9-sgRNA expression vectors for plant transformation (Figure 5a; Table S6). To generate either in-frame or out-of-frame αKNL2s mutants, several constructs were designed. Each construct contained two sgRNAs targeting the same gene and therefore was able to generate large fragment deletions ranging from 69 bp to 1648 bp (Figure 5b, c; Table S3). All constructs were delivered into cowpea cells using an *Agrobacterium*-mediated embryonic axis-based transformation protocol previously published (Che et al. 2021). Transgenic T0 regenerants were selected on medium containing Spectinomycin, with an overall transformation efficiency of 15-20% (Figure 6a). PCR confirmed the presence of the transgenic construct with construct-specific primers and green fluorescence emission of tGFP in transgenic plants (Figure 6b, c). All independent lines obtained (more than 50 primary transformants per construct) were transferred to the greenhouse, and T1 seeds were collected. PCR analysis of T1 transgenic seedlings with *αKNL2.1*/*αKNL2.2* gene-specific primers revealed the presence of large expected deletions within the targeted genes (Figure S4). Subsequent sequencing analysis of respective amplicons of genomic DNA confirmed the presence of expected deletions within individual or both genes at the expected positions defined by sgRNA (Figure S5).

**Figure 5.**
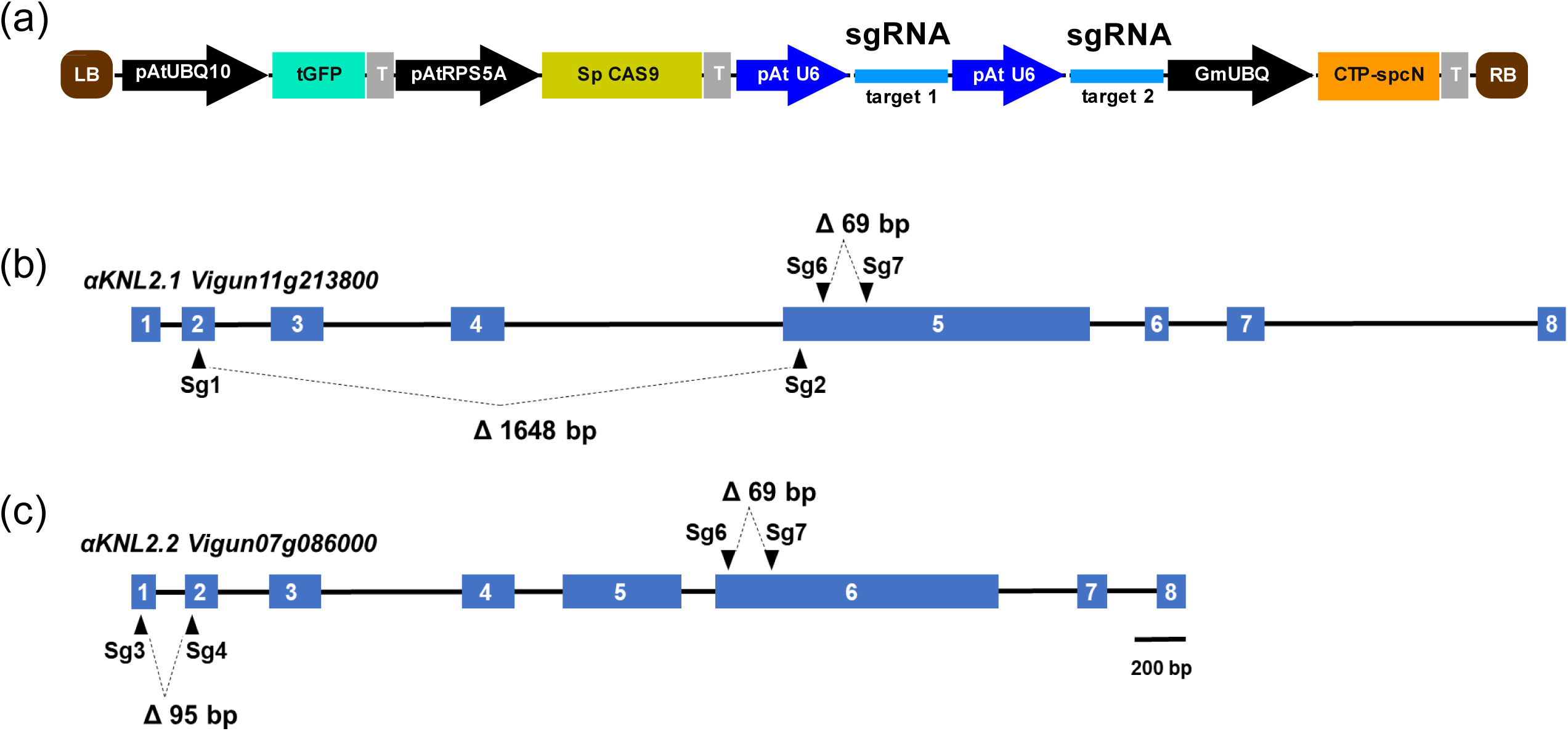
CRISPR/Cas9-mediated gene editing of αKNL2s in cowpea. (a) Schematic representation of the constructs used in this study. Each construct contains two sgRNAs for targeting the genomic sequence either αKNL2.1 or αKNL2.2. pAtUBQ10, Arabidopsis ubiquitin-10 gene promotor; tGFP, turboGFP (reporter); pAtRPS5A, Arabidopsis ribosomal protein 5a promoter; Sp CAS9, Streptococcus pyogenes CAS9; pAt U6, Arabidopsis U6 promoter; sgRNA, guide RNA; GmUBQ, Glycine max polyubiquitin promoter; CTP-spcN, Streptomyces spectabilis spectinomycin resistance gene (selectable marker); LB and RB, left and right T-DNA borders. (b, c) αKNL2 genes structure and target sequence locations. Blue boxes and the black lines indicate exons and introns, respectively. Scale bar =200 (bp) base pairs. Positions of sgRNA in targeted genes are shown.

**Figure 6.**
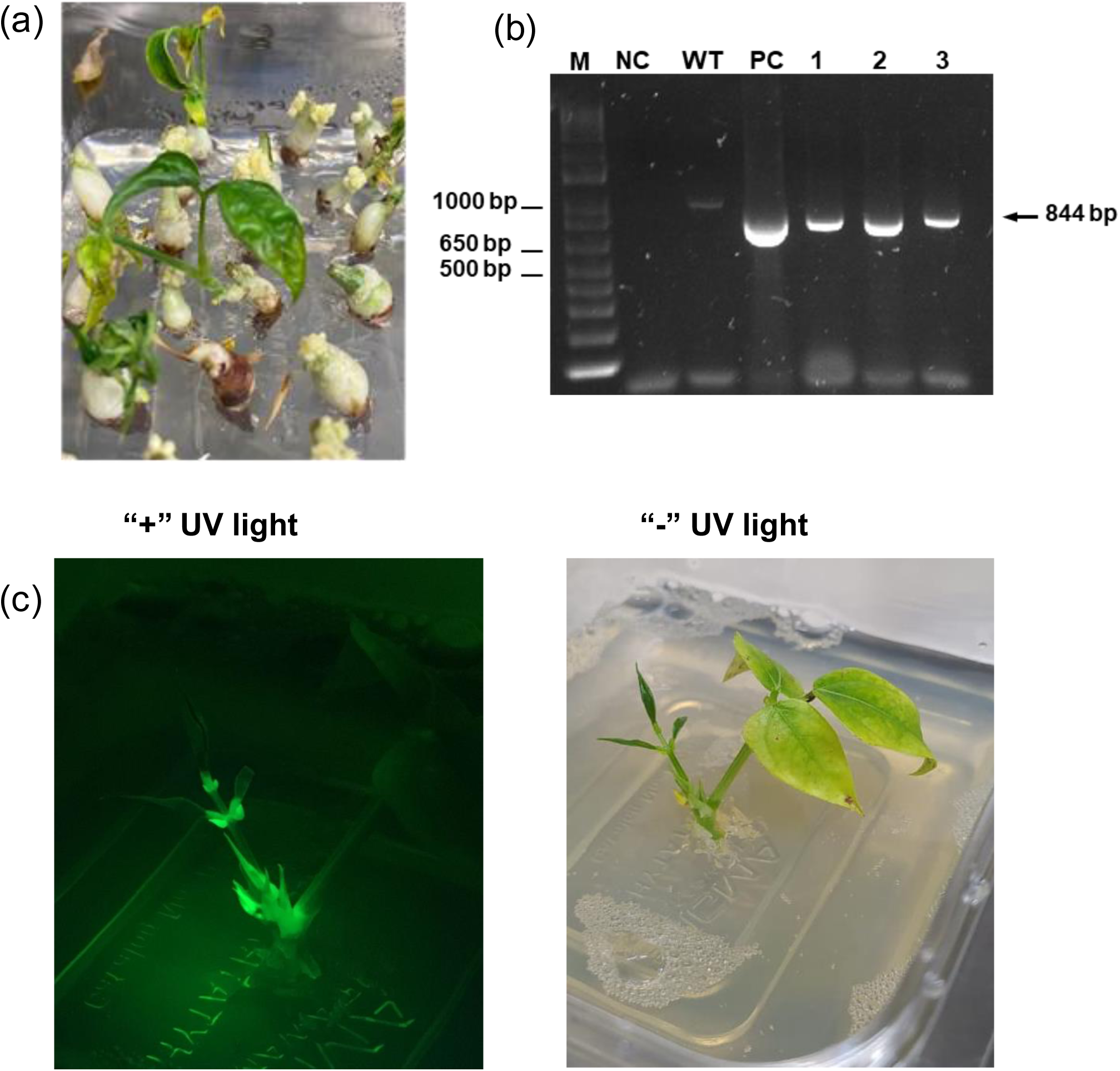
Selection and analysis of transgenic cowpea plants expressing the CRISPR/Cas construct. (a) Cowpea explants with transgenic shoots on spectinomycin-containing medium. (b) PCR amplification with spectinomycin resistance gene-specific primers. M, molecular size marker; NC, water and PC, isolated plasmid DNA used as negative control and positive control, respectively; WT, non-transgenic wild-type *V. unguiculata* cv. IT86D-1010; 1-3, the independent transgenic cowpea lines. Arrow indicates the amplified spcN fragment (844 bp). (c) (c) Transgenic cowpea plantlet under GFP filter and bright field.

Moreover, all T1 seedlings were screened for tGFP fluorescence, and their genomic DNA was assayed with Cas9 gene-specific primers. Such combined analysis allowed the identification of plants that lacked a transformation cassette (Cas9/tGFP null segregants) (Figure S6). Ten independent lines for each gene-editing experiment, that were identified as transgene-free cowpea CRISPR mutants, were transferred to the greenhouse and subjected to one round of crossing with wild-type plants. The resulting F1 generation was selfed to produce homozygous mutant lines. Finally, a set of *αknl2* mutants was generated. We produced single homozygous *αknl2* out-of frame (of) knock-out mutants for each of individual *αKNL2* gene (αknl2.1-of αknl2.1-of/αKNL2.2 αKNL2.2 and αKNL2.1 αKNL2.1/αknl2.2-of αknl2.2-of), a double homozygous in-frame (if) mutant expressing truncated version of both genes (αknl2.1 - if αknl2.1-if/αknl2.2-if αknl2.2-if), and a mutant which was homozygous for αknl2.2-of and heterozygotes for αknl2.1-of (αknl2.1-ko αKNL2.1/αknl2.2-of αknl2.2-of) (Table S4). All generated mutants displayed normal growth and did not show any other plant developmental abnormalities (Figure S7). Unfortunately, our multiple attempts to generate homozygous double-mutant knock-out plants were unsuccessful. It is worth noting that plants possessing αknl2.1-of αKNL2.1/αknl2.2-of αknl2.2-of genotype can be easily generated via crossing of the corresponding single-gene mutants and subsequent selfing. However, further screening of the progeny after self-fertilization of such plants still did not yield any homozygous double mutants. Visual inspection of the pods from plants after selfing revealed that each pod, in addition to “normal” seeds, also contained seeds with an abnormal phenotype (small-sized “aborted” seeds) (Figure S8). This observation led us to conclude that homozygous double knock-out mutants were most likely not physiologically viable. Therefore, producing such mutants could be very challenging, or even impossible.

### Evaluation of αKNL2s and CENH3.1 levels in CRISPR mutants

Finally, a comparison between wild-type and various αKNL2 mutant lines was performed by immunostaining mitotic root meristems with anti-αKNL2 and anti-CENH3.1 antibodies (Figure S9). After indirect immunostaining with anti-αKNL2, a bright fluorescence signal was observed in the centromeric region, which was confirmed by colocalization with antibodies against cowpea CENH3. The presence of such a signal in single KO mutants (αknl2.1 or αknl2.2) can be explained by the availability of a second non-mutated isoform of αKNL2 protein. At the same time, the appearance of centromeric anti-KNL2 signals in the αknl2.1-if αknl2.1-if/αknl2.2-i f αknl2.2-if mutant suggests that the αknl2.1-if and αknl2.2-if mutant forms of the proteins (truncated proteins) can retain activity and serve the same function as the wild-type αKNL2 proteins. No decrease of anti-αKNL2 or CENH3 signals in the nuclei of all tested αknl2 mutants compared to the wild type was detected.

### Analysis of the haploid induction ability of cowpea αKNL2 mutants

To determine whether mutations in αKNL2 genes could trigger *in vivo* haploid induction in cowpea, as demonstrated for *A. thaliana* (Ahmadli et al. 2023), we used the generated single- and double-mutants as mother and father parents in crosses with wild type under controlled greenhouse conditions. The ploidy level of F1 progeny from such crosses was determined by flow cytometry. Unfortunately, among the 100 F1 plants analyzed, none was found to be haploid (Figure S10). These results suggest that knocking out αKNL2 genes does not induce *in vivo* haploid induction in cowpea at a high frequency.

## Discussion

The architecture and function of the plant kinetochore have been investigated, and progress has been made in characterizing a number of genes encoding structural and regulatory kinetochore proteins. Acquired knowledge substantially broadened our understanding of the processes of kinetochore assembly, directed chromosome movements and congression, and spindle assembly checkpoint signaling (Lara-Gonzalez et al. 2021; Naish and Henderson 2024; Kozgunova 2025; Xie et al. 2024).

αKNL2 protein is suggested to be one of the centromere-licensing factors responsible for CENH3 loading at centromeres and promoting kinetochore assembly (Lermontova et al. 2013; Yalagapati et al. 2025). The presence of the CENPC-k motif characterizes the cowpea αKNL2 homologs identified in this study, in addition to the conserved SANTA domain. Therefore, according to the classification by (Zuo et al. 2022), both can be defined as αKNL2 proteins. While *A. thaliana* has only one *αKNL2* gene, diploid cowpea possesses two paralogous genes, αKNL2.1 and αKNL2.2, each encoding distinct isoforms that may perform specific functions in addition to housekeeping functions. The existence of either two or three isoforms of αKNL2 has been reported in the polyploids *Brassica napus, B. juncea, Lepidium meyenii* and *Camelina sativa* (Zuo et al. 2022). However, data on their specific function in mitosis and meiosis remain obscure. Notably, cowpea similarly encodes two functional CENH3 variants (Ishii et al. 2020). Whether a functional relationship exists between the duplication of these two genes remains to be elucidated.

*αKNL2.1* was highly expressed in all tissues examined, including root meristems and apical meristems of leaves, where cells demonstrated high mitotic activity. Therefore, we assume that αKNL2.1 is the primary protein responsible for centromere licensing and CENH3 loading. Since expression of αKNL2.2 was also relatively high in all tissues analyzed, the presence of αKNL2.2 is likely necessary for cooperative work with αKNL2.1 in the process of kinetochore formation. However, under certain conditions, the αKNL2.2 may have a leading role in CENH3 loading. Our analysis of generated CRISPR/Cas9 knock-out mutants corroborates this assumption.

Indeed, our fluorescent protein tagging and immune assays demonstrated co-localization of both αKNL2 isoforms at centromeres. This suggests that both variants are involved in CENH3 loading and/or stabilization of incorporated CENH3 into centromeric nucleosomes. In line with the previously reported timeline of αKNL2 centromere association in *A. thaliana* (Lermontova et al. 2013) and the moss *P*. *patens* (Kozgunova et al. 2019), both cowpea αKNL2 variants colocalize with interphase centromeres, but in mitotic cells, αKNL2 dissociates from centromeres from prophase to anaphase. Thus, there is similarity in αKNL2 regulation between mosses and flowering plants.

Generated *αknl2.1* and *αknl2.2* cowpea single knock-out mutants did not exhibit any phenotypic alterations compared with the wild-type plants. This is in contrast with previous findings in *A. thaliana*, where the *αknl2* T-DNA insertion mutant plants were characterized by abnormal growth and leaf shape, multiple shoot formation and reduced size of siliques (Lermontova et al. 2013). Such differences can be explained by the fact that *A. thaliana* encodes only one *αKNL2* gene, and its knock-out results in complete depletion of αKNL2.

Our data indicate that αKNL2.1 and αKNL2.2 are not individually required for vegetative or floral development in cowpea, pointing to functional redundancy between these isoforms. Thus, the presence of a single α-isoform is sufficient to ensure the loading of CENH3 to centromeric chromatin and for the normal process of progression of kinetochore formation. At the same time, we did not obtain double-homozygous knock-out mutants of α*KNL2,* suggesting that α*KNL2* plays an essential role during early seed development. A high level of seed abortion (up to 30%) was also reported before for *A. thaliana αknl2* knock-out mutant (Lermontova et al. 2013). Knock-out of genes encoding proteins involved in chromosome segregation and cell division very often leads to the arrest of seed development. In particular, seed abortion was reported for several kinetochore protein null mutants, including *mis12* (Sato et al. 2005), *cenh3* (Ravi et al. 2010), *nnf1* (Allipra et al. 2022), and *ttn9* (Zhang et al. 2025).

In our study, attempts to generate haploids in cowpea using single- and double αknl2 mutants were unsuccessful. However, while the single- and double *αknl2* mutants failed to induce haploidy in cowpea, the limited sample size per genotype suggests these results are preliminary. Further testing with larger populations may be required to fully assess their potential. In *A. thaliana*, the *αknl2* mutant yielded a haploid induction rate of 1% under standard growth conditions (Ahmadli et al. 2023). Moreover, applying short-term temperature stress to *αknl2* cowpea mutants, a technique proven effective in *A. thaliana* (Ahmadli et al. 2023), might improve induction rates. Also, the application of *βknl2* mutants alone or in combination with *αknl2* mutations might allow the induction of haploids in cowpea. Alternatively, uniparental gametic depletion of the KNL2 protein or other kinetochore proteins, as recently demonstrated for *Arabidopsis* CENH3 (Somasundaram et al. 2026), might represent a viable strategy to induce haploids in cowpea.

## MATERIALS AND METHODS

### Plant materials and growth conditions

Seeds of wild-type *V. unguiculata* cv. IT86D-1010 and CRISPR mutants generated on the same genetic background were germinated on wet filter paper and afterwards transferred to the 20 cm-diameter pots containing commercial soil substrate. Plants were grown in a greenhouse at 26°C/18°C day/night temperature, with 60% humidity and 16 h light per day photoperiod. The pots were watered twice a day.

### Phylogenetic tree

*Fabaceae* species sequences of KNL2 were retrieved from the Gramene (https://plants.ensembl.org; (Tello-Ruiz et al. 2022)) and Phytozome (https://phytozome-next.jgi.doe.gov; (Goodstein et al. 2012)) databases. The phylogenetic tree was constructed in MEGA 11 (Tamura et al. 2021)) using the Maximum likelihood method with 1000 bootstrap replicates.

### RNA extraction and RT-qPCR assays

Tissue samples of roots, leaves, stems, flowers, pods, and seeds were collected in three biological replicates per sample. Total RNA was extracted using Trizol™ Reagent (Invitrogen, Thermo Fisher Scientific, Germany). After treatment by Turbo DNA free (Thermo Fisher Scientific, Waltham, MA, USA), the total RNA was reverse transcribed into cDNA using RevertAid H Minus First Strand cDNA Synthesis Kit (Thermo Fisher Scientific Baltics, Lithuania).

Expression levels of the *αKNL2.1* and *αKNL2.2* genes across different organs of cowpea were assayed by quantitative RT-PCR with PowerUp SYBR Green Master Mix and QuantStudio 6 Flex system (Applied Biosystems, Thermo Fisher Scientific). Amplification conditions were 2 min at 50°C, 10 min at 95°C, 40 cycles each of 15 s at 95°C followed by 1 min at 60°C; 15 s at 95°C; 20 s at 60°C; and 15 s at 95°C. Two house-keeping genes actin (Vigun04g203000) and ubiquitin-conjugating enzyme E2 (Vigun07g012200) were used as internal references for the quantification of gene expression levels. The sequences of gene-specific primers used are listed in Table S5.

### Design and *in vitro* cleavage assay of guide RNAs

The CRISPR-P 2.0 online tool (Lei et al. 2014) was used to design a set of guide RNAs targeting the αKNL2.1/αKNL2.2 genes in cowpea. *In vitro* efficiency analysis of different guide RNAs was performed using the RNP complexes containing commercial *Streptococcus pyogenes* Cas9 endonuclease (New England Biolabs, Frankfurt am Main, Germany) and synthetic gRNAs (Integrated DNA Technologies, Inc.,Commercial Park Coralville, USA). To prepare sgRNA, synthetic tracrRNA and crRNA were mixed in equimolar concentrations to get a final duplex concentration of 10 μM. Afterwards, samples were heated to 95°C for 5 min, followed by slow cooling to room temperature. The RNP complexes were assembled by mixing sgRNA (70 nM) and Cas9 (70 nM) in NEB buffer 3.1 (New England Biolabs, Germany) containing 50 mM Tris-HCl pH 7.9, 100 mM NaCl, 10 mM MgCl2, and 100 µg/ml BSA, and incubating for 10 min at 25 °C. Afterwards linearized plasmid bearing genomic DNA sequence of *αKNL2.1/αKNL2.2* (7 nM) was added, and mixtures were incubated at 37°C for 1h. To stop the reaction, Invitrogen Proteinase K (Thermo Fisher Scientific, Germany) was added, and the samples were incubated for 10 min at 56°C. Cleaved products were analyzed by 0.8% agarose gel electrophoresis.

### Plasmid construction

All plasmids were generated using Golden Gate modular cloning technology and MoClo Toolkit (vector toolkit; (Weber et al. 2011; Engler et al. 2014). Individual parts: AtpRPS5A, CAS9, At act2 ter, AtUBQ10, OCS ter, pαKNL2.1pro, pαKNL2.2pro GUS, 35S ter, CAMV35Spro, CDS_αKNL2.1, CDS_αKNL2.2, and EYFP were first amplified with primers containing 5’ extensions with BpiI restriction sites (Table S5) and then subsequently cloned into respective level 0 destination vectors using the Golden-Gate restriction-ligation protocol published before (Grutzner and Marillonnet 2020). Afterwards, the separate modules were assembled into complete transcription units in MoClo level-1 cloning vectors using *Bsa*I enzyme (Table S6).

To generate a set of guide RNAs under AtU6 promoter, the pICH86966:: AtU6p::sgRNA PDS (Addgene plasmid 46966) was used as a template for amplification with specific guide RNA primers (Table S5). Subsequently, PCR products were cloned together with the pICSL01009::AtU6p plasmid (Addgene 46968)) via a BsaI Golden Gate cloning reaction either into pICH47751 or pICH47761 level-1 acceptor vectors (Table S6). Finally, level-1 modules were assembled into the binary vector pAGM4723 to obtain CRISPR/Cas expression, EYFP-CDS, and promoter GUS-fusion constructs (Figure 5, Table S6) using the *Bpi*I restriction enzyme. All generated constructs were verified by whole plasmid sequencing (Eurofins Genomics, Ebersberg Germany) and introduced into the *Agrobacterium tumefaciens* via electroporation.

### αKNL2 gene editing in *V. unguiculata*

For the transformation, the thymidine-auxotrophic *Agrobacterium tumefaciens* strain (LBA4404Thy-) (Ranch et al. 2010) carrying the binary vector pAGM4723 with two gene-specific sgRNA, Cas9, tGFP (visual marker) and spcN (selectable marker for spectinomycin resistance) was used. The transformation procedure for the embryonic axis of cowpea explants was conducted as described previously (Che et al. 2021). Regeneration and selection of primary transformants were achieved on MS medium (Murashige and Skoog 1962) containing 50 mg/l spectinomycin, 15 mg/l meropenem, BAP 0.5 mg/l, kinetin 0.5 mg/l, 3 % sucrose, and 0.8 % agar for solidification at pH of 5.6. After rooting on MS medium with IBA 0.1 mg/l, T0 plants were transferred to the soil and allowed to set seeds in the greenhouse. The green fluorescence of tGFP in T0 and T1 seedling tissues was screened using a NIGHTSEA fluorescent lamp with an RB-Green filter (440–460 nm) and an MZ10 F Modular Stereo Microscope (Leica, Germany), respectively.

Genomic DNA from young leaf tissue was extracted according to the standard CTAB method (Doyle and Doyle 1990). PCR amplification with OneTaq DNA Polymerase (NEB, Germany) and construct-specific primers listed in Supplementary Table S5 was carried out according to the manufacturer’s instructions, to reveal integration of the T-DNA of the binary vectors into genomic DNA of the primary transformants. For the analysis of generated mutations, the PCR amplification with *αKNL2.1/ αKNL2.2* gene-specific primers (Table S5) encompassing CRISPR mutagenic sites was employed. The resulting PCR products were cloned into the pCR8/GW/TOPO vector (Invitrogen, Thermo Fisher Scientific, Germany) and sequenced.

### GUS Staining

β-glucuronidase enzyme activity was detected histochemically according to (Jefferson et al. 1987)). In brief, samples were incubated overnight at 37°C in 100 mM sodium phosphate buffer containing 5-bromo-4-chloro-3-indolyl-beta-D-glucoroni de (X-Gluc, Duchefa Biochemie, Haarlem, The Netherlands) as a substrate. After incubation, the samples were washed with 70% ethanol to remove chlorophyll, and the stained plant material was photographed.

### Transient expression and indirect immunostaining

*A. tumefaciens strain* GV2260 harbouring one of the following binary vectors: 35S::eYFP::αKNL2.1-pAGM4723, 35S::eYFP::αKNL2.2-pAGM4723, and p35S::p19 was used to infiltrate leaves of four-week-old cowpea plants. Infiltration into the abaxial side of leaves was performed using a needleless 1 mL syringe as described previously (Leite Dias et al. 2025). The infiltrated leaves were harvested on the fourth day after agroinfiltration, and tissues were fixed in Tris-buffer (10mM Tris, 100 mM NaCl, 10 mM EDTA, 0.1% Triton X-100, pH7.5) containing 4% paraformaldehyde (PFA) initially on ice under vacuum for 5 minutes followed by 30 minutes without vacuum. Afterwards, samples were washed twice for 3 minutes with cold buffer without PFA. To isolate the nuclei, the fixed material was chopped in buffer composed of 15 mM Tris, 2 mM EDTA, 0,5mM spermine, 80 mM KCI, 20 mM NaCl, 15 mM 2-Mercaptoethanol, 0,1% Triton X-100 at pH7.0. The nuclei suspension was filtered through a 50 µm mesh.

The filtered suspension was transferred to glass slides using a Cytospin 3 cytocentrifuge (Thermo Scientific, Cheshire, United Kingdom). The slides were washed twice with 1x -buffered saline (PBS) (Sigma-Aldrich, St. Louis, USA) and then blocked with 4% bovine serum albumin (BSA) for 1 hour. As the primary antibodies, the rabbit anti-CENH3 antibody reported previously (Ishii et al., 2020) and the rabbit anti-αKNL2 peptide antibody (LifeTein LLC, Hillsborough, USA) produced for the current study were used. For immunofluorescence staining, secondary rhodamine-conjugated goat anti-rabbit IgG (Diagnostica Vertrieb GmbH, Hamburg, Germany) were used. Immunostained leaf nuclei were examined using fluorescence microscope Olympus BX61 (Olympus, Tokyo, Japan). Photographs were taken using an ORCA-ER CCD camera (Hamamatsu, Japan).

### Flow cytometry

The ploidy analysis of nuclei isolated from young leaves of cowpea plants was performed using BD FACSAria IIu cell sorter equipped with 375 nm near-UV laser and 488 nm blue laser (BD Biosciences, San Jose, CA, USA) as described in detail before (Somasundaram et al. 2026).

## Acknowledgements

We thank Sylvia Swetik and Sylvane Stegmann (IPK) for technical assistance and Inna Lermontova (IPK) for discussion. This work was supported in part by a sub-award from the University of Queensland to Leibniz Institute of Plant Genetics and Crop Plant Research Gatersleben (IPK) under the Hy-Gain project (INV-002955) from the Gates Foundation. The conclusions and opinions expressed in this work are those of the author(s) alone and shall not be attributed to the Foundation. Under the grant conditions of the Foundation, a Creative Commons Attribution 4.0 License has already been assigned to the Author Accepted Manuscript version that might arise from this submission.

## CONFLICT OF INTEREST

The authors declare no conflicts of interest. This article does not contain any studies with human or animal participants.

## Authors contributions

A.K., J.-P.V.-C. and A.H. designed the experiments. A.K., I.A.-M., G.L-M., J.L., O. R.- M. and J.F. performed the experiments. A.K., J.-P.V.-C and A.H. wrote the paper.

## Supplementary Figures

**Figure S1.**
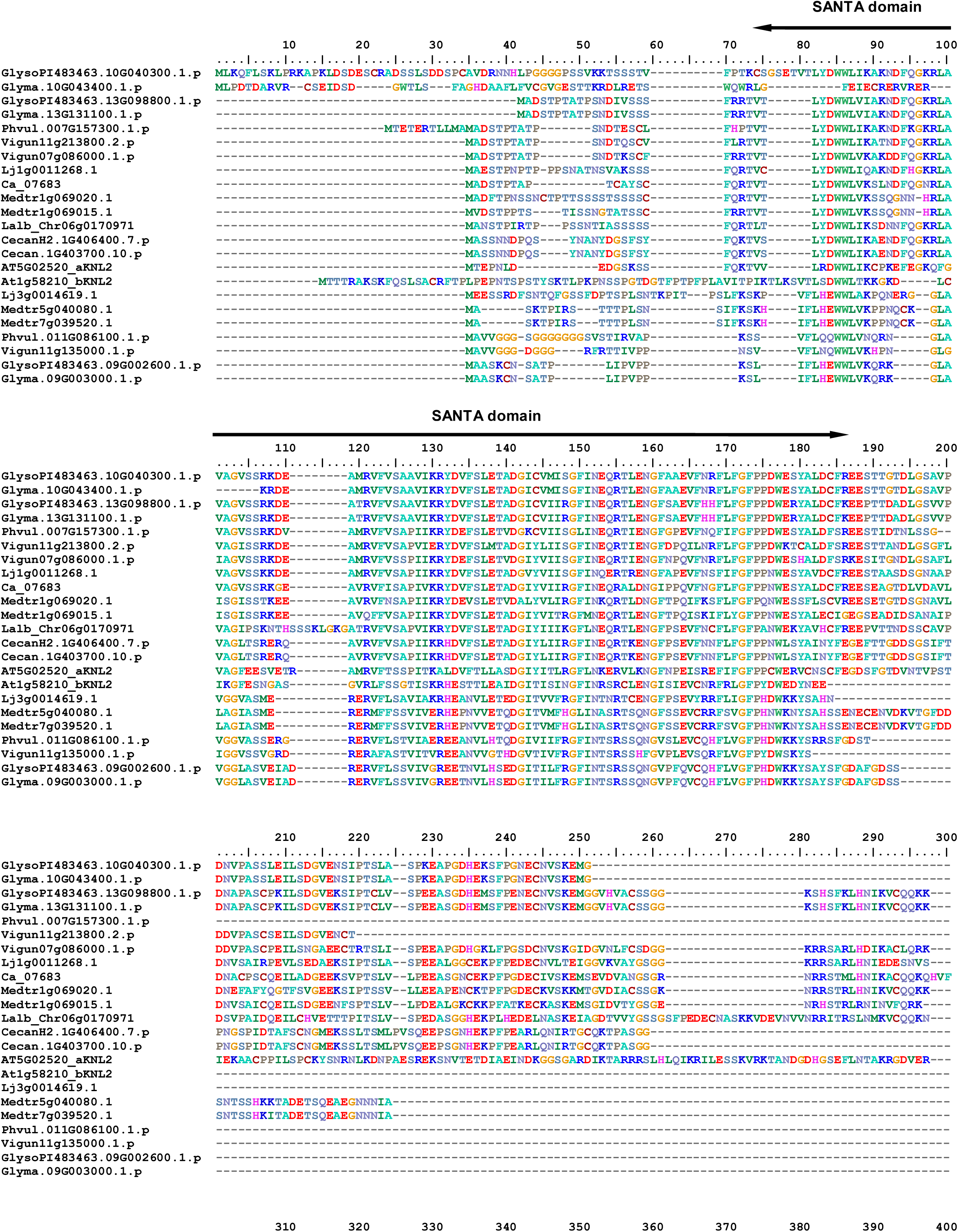
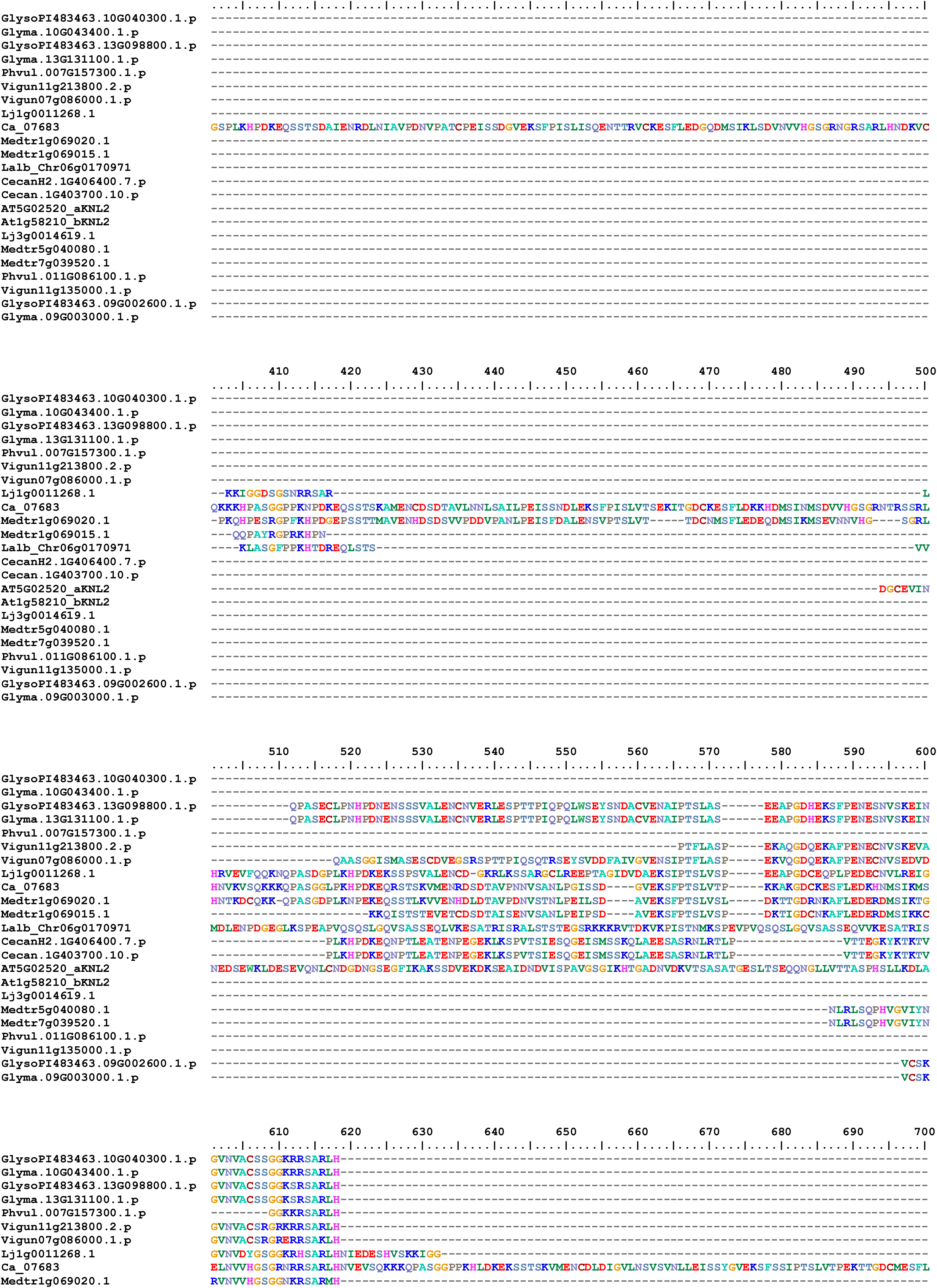
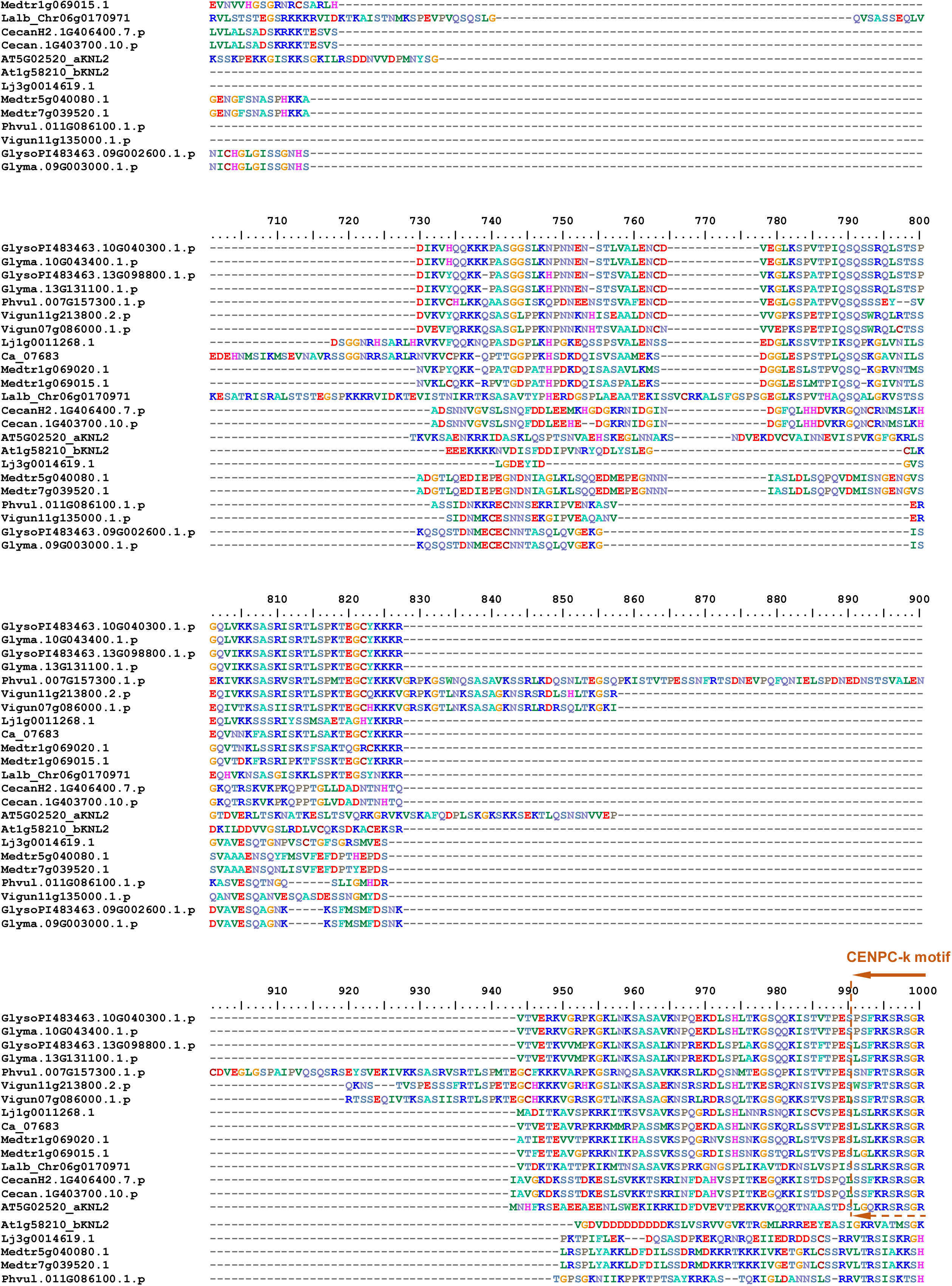
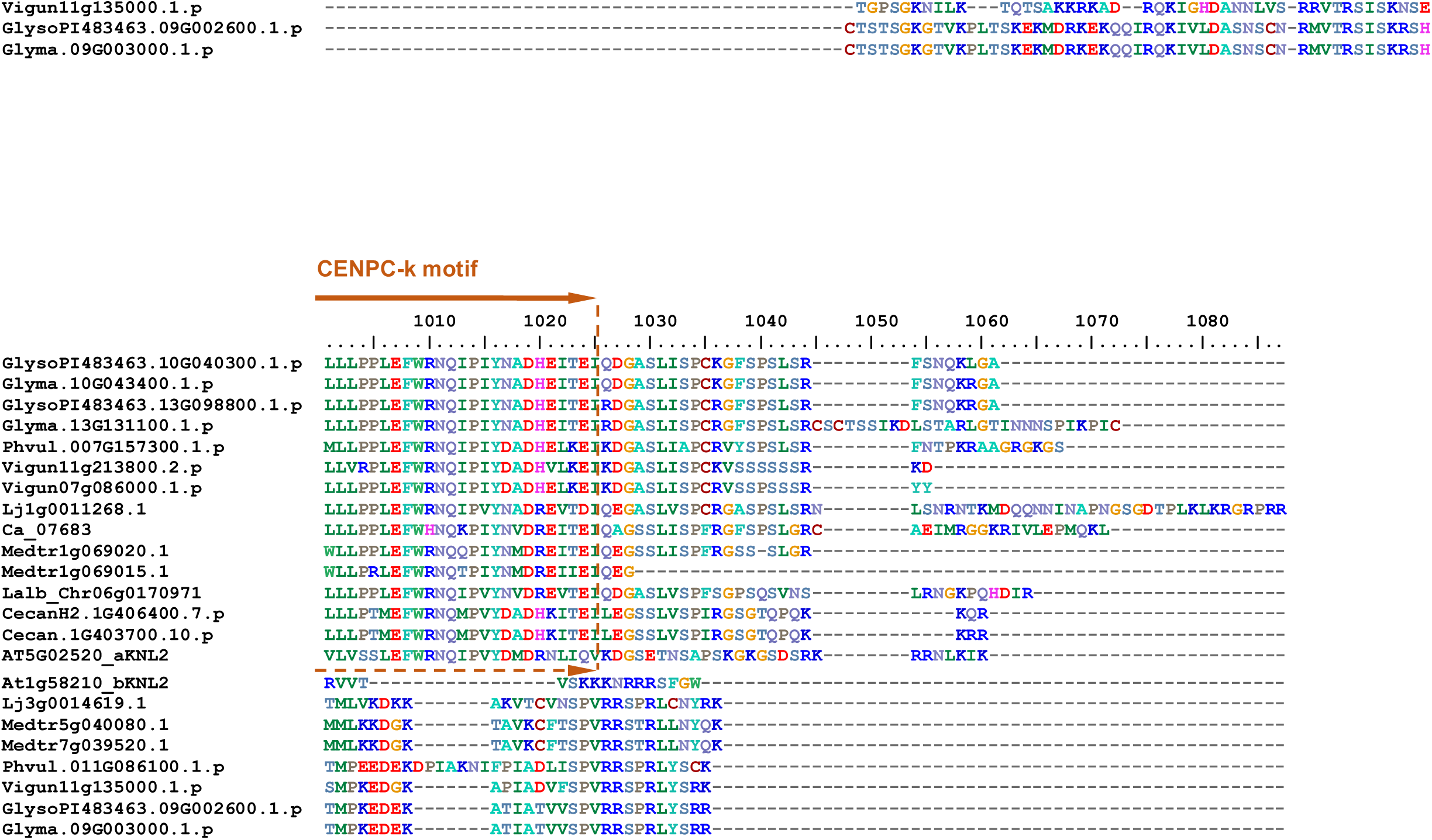
Amino acid sequence alignment of Fabaceae αKNL2 proteins. The conserved SANTA domain and typical CENPC-k motif of αKNL2 are indicated.

**Figure S2.**
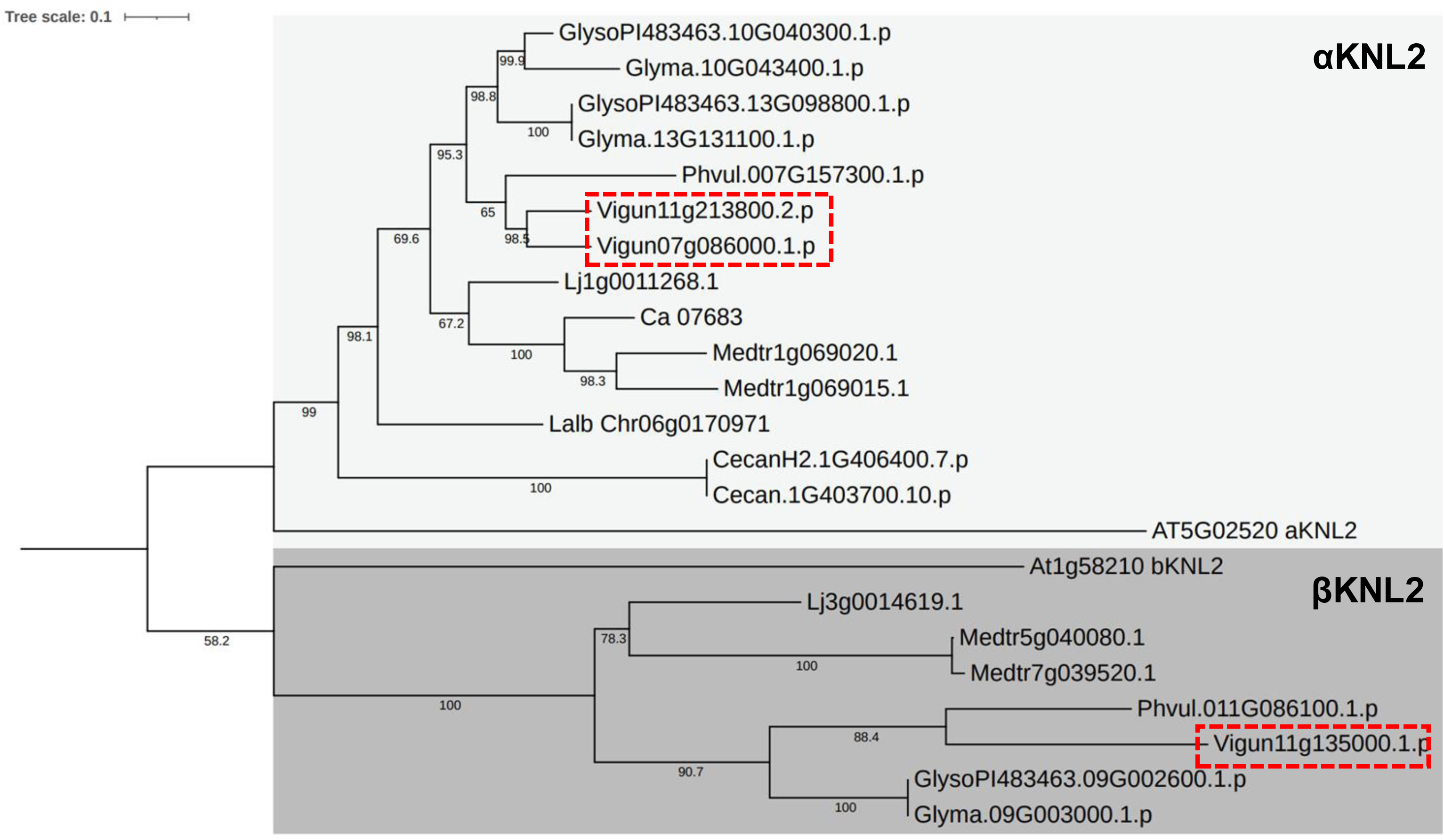
Maximum likelihood phylogenetic tree of KNL2 homologs of the family Fabaceae. Phylogenetic analysis was carried with protein sequences from *Glycine soja* (GlysoPI483463.10G040300.1.p; GlysoPI483463.13G098800.1.p; GlysoPI483463.09G002600.1.p), *Glycine max* (Glyma.10G043400.1.p; Glyma.13G131100.1.p; Glyma.09G003000.1.p), *Phaseolus vulgaris* (Phvul.007G157300.1.p; Phvul.011G086100.1.p), *Vigna unguiculate* (Vigun11g213800.2.p cowpea αKNL2.1; Vigun07g086000.1.p αKNL2.2; Vigun11g135000.1.p cowpea βKNL2), *Lotus japonicus* (Lj1g0011268.1; Lj3g0014619.1), *Cicer arietinum* (Ca_07683), *Medicago truncatula* (Medtr1g069020.1; Medtr1g069015.1; Medtr5g040080.1; Medtr7g039520.1*), Lupinus albus* (Lalb_Chr06g0170971), *Cercis canadensis* (CecanH2.1G406400.7.p; Cecan.1G403700.10.p), and *A. thaliana* (AT5G02520 αKNL2; At1g58210 βKNL2). Calculated bootstrap values are shown near corresponding nodes. The scale at the top left corner represents the number of substitution events (10 per 100 amino acids).

**Figure S3.**
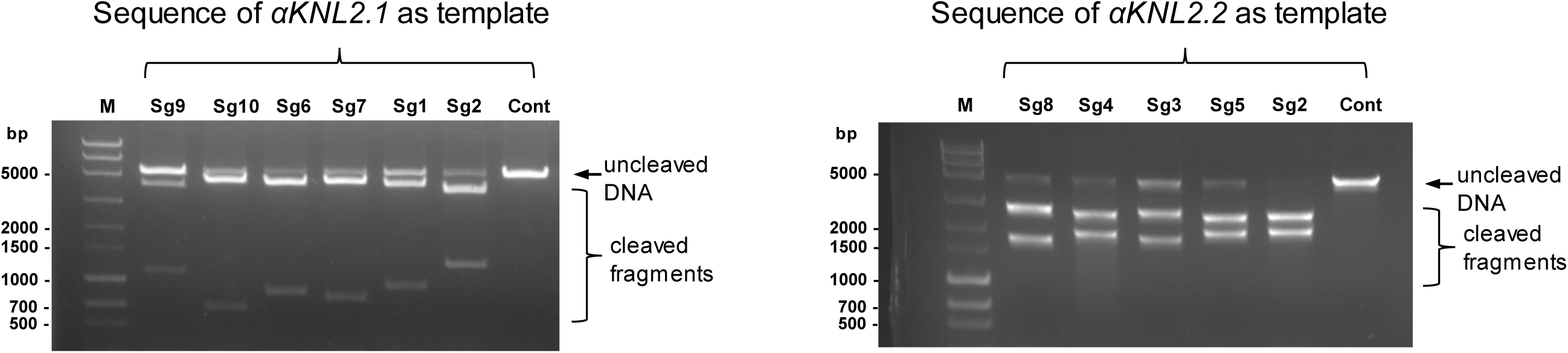
Exploring the ability of designed sgRNAs to target αKNL2 sequences by means of *in vitro* cleavage assay. M, molecular size marker MassRuler Express Forward DNA Ladder Mix (Thermo Scientific); Sg1-Sg10, targeted gene region digested with Cas9 and respective sgRNA; Cont, negative control that lacked sgRNA that was included for comparison.

**Figure S4.**
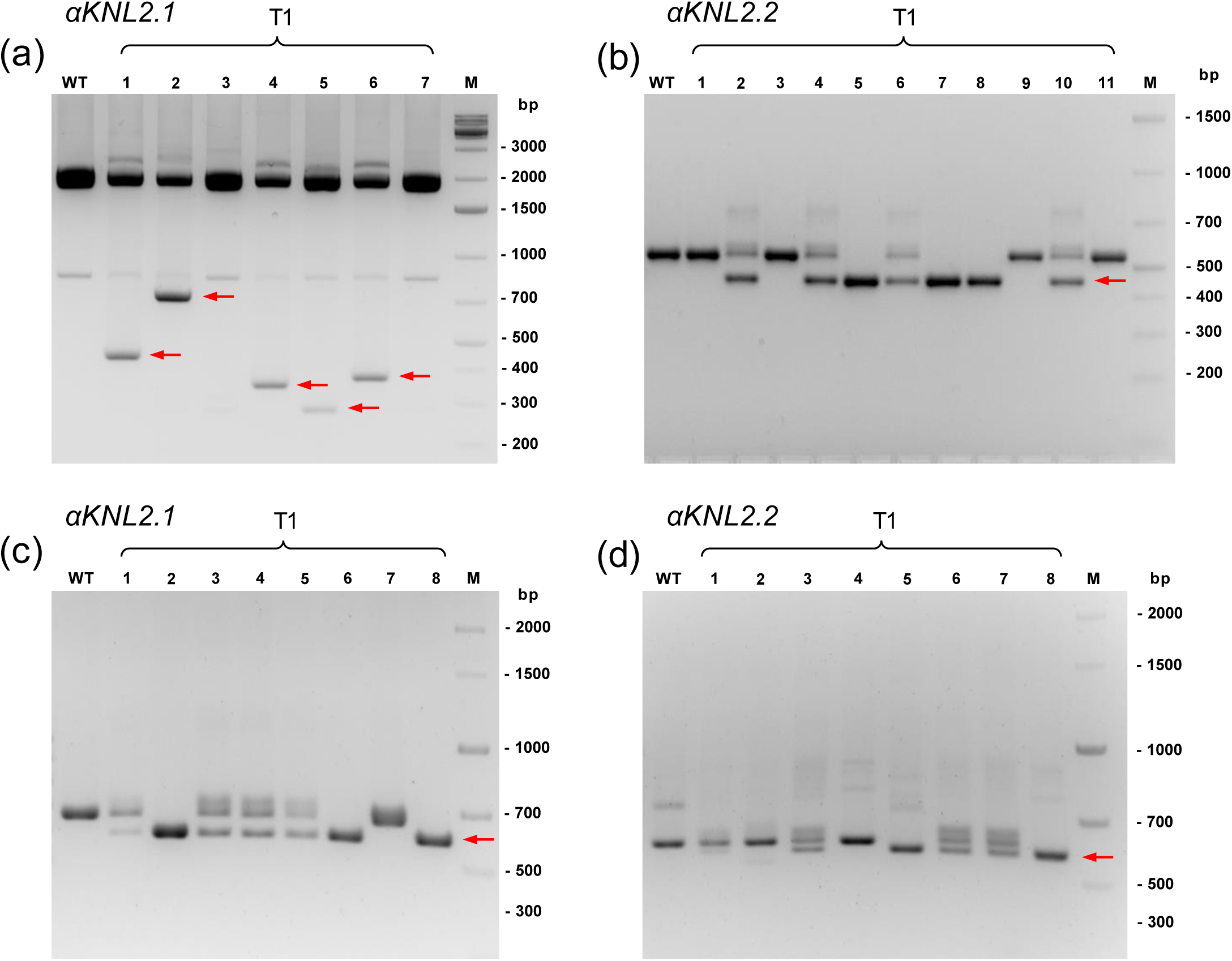
Detection of targeted deletions (out-of-frame knockout; a, b) and (in-frame mutations; c, d) using PCR with gene-specific primers. M, DNA marker; lanes 2–9, PCR products of genomic DNA with gene specific primers. M, DNA marker; lanes 1-11, PCR products of genomic DNA from independent CRISPR/Cas-mutagenized T1 lines. Deletions are shown by red arrows.

**Figure S5.**
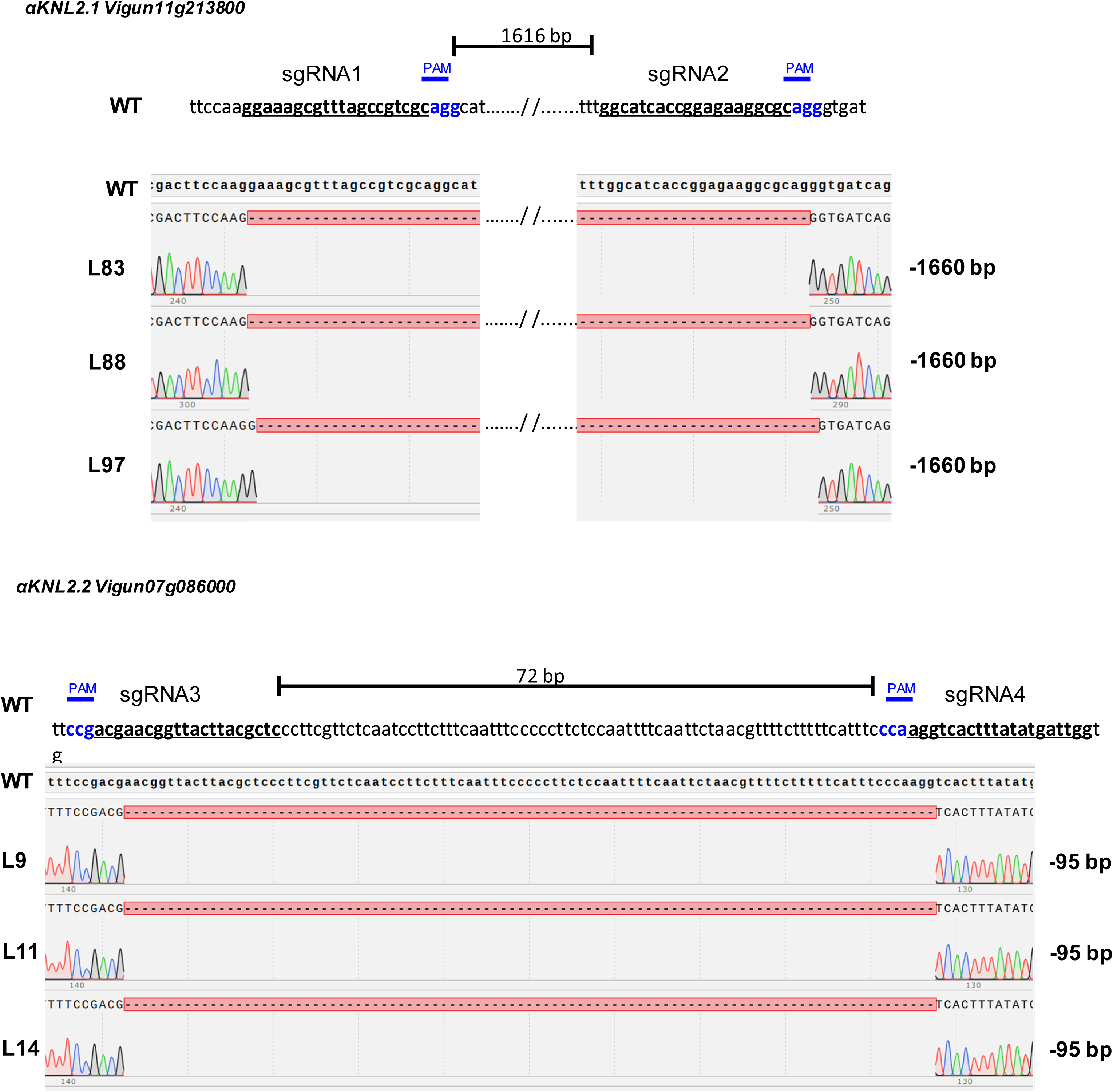
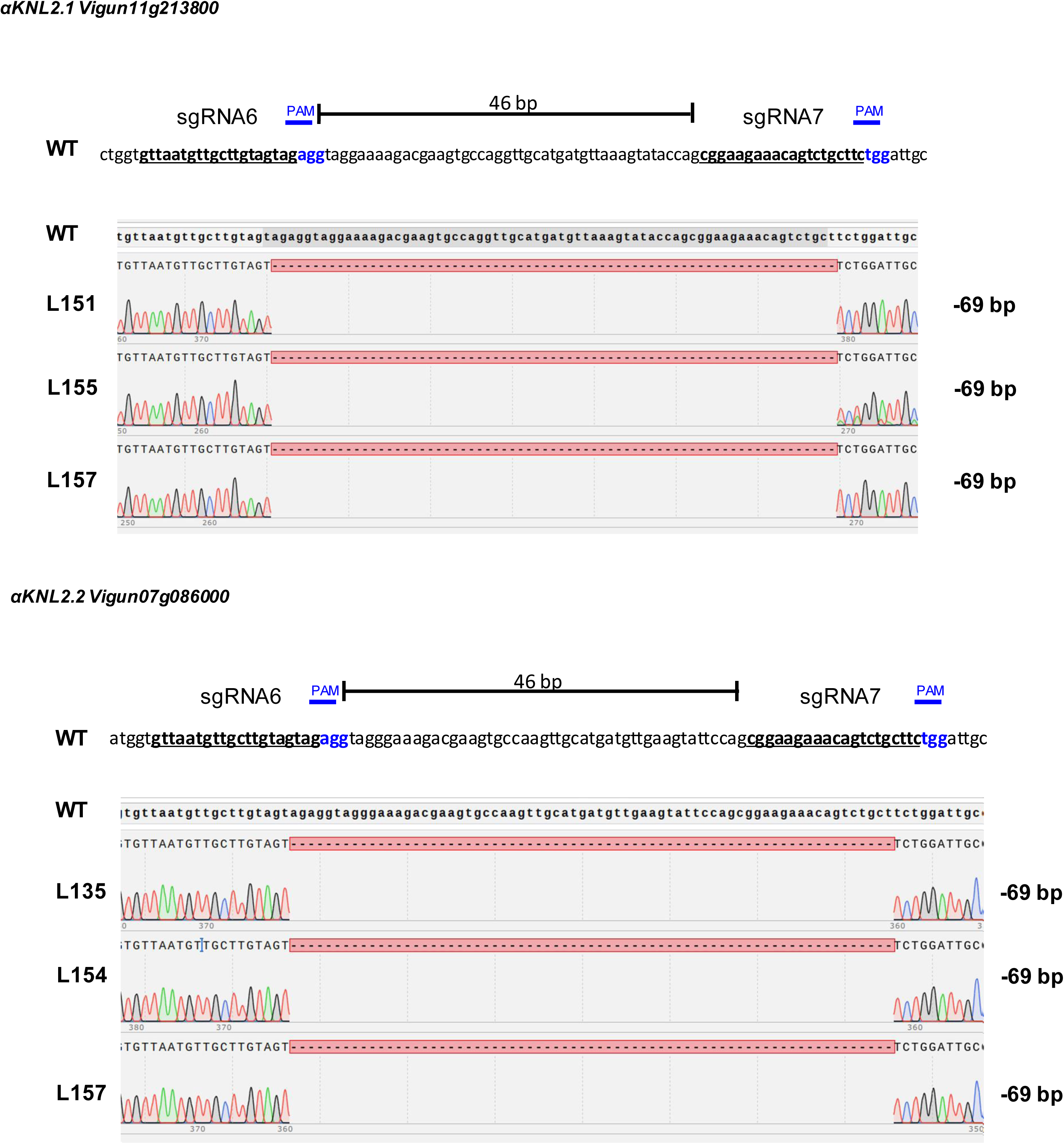
Representative chromatograms of αKNL2 sequences with generated nucleotide deletions. Sequences corresponding to the sgRNA and PAM are highlighted.

**Figure S6.**
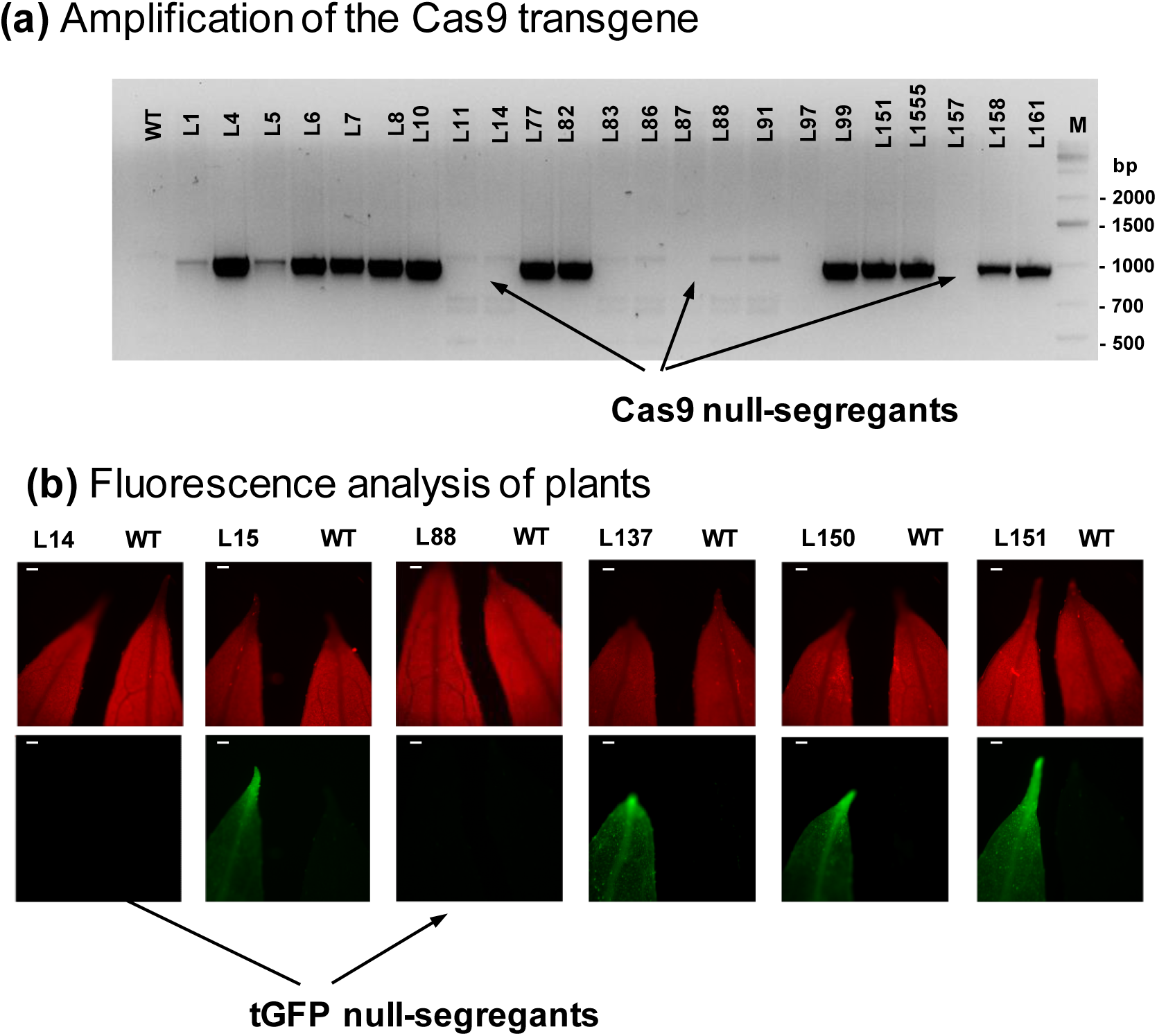
Search for Cas9/tGFP null segregants. (a) Amplification of gDNA of various T1 lines with Cas9 specific primers. (b) Fluorescence analysis of T1 plants. Red is autofluorescence of chlorophyll. Green is tGFP fluorescence signal observed in leaf of T1 plant. The scale bars correspond to 200 µm .

**Figure S7.**
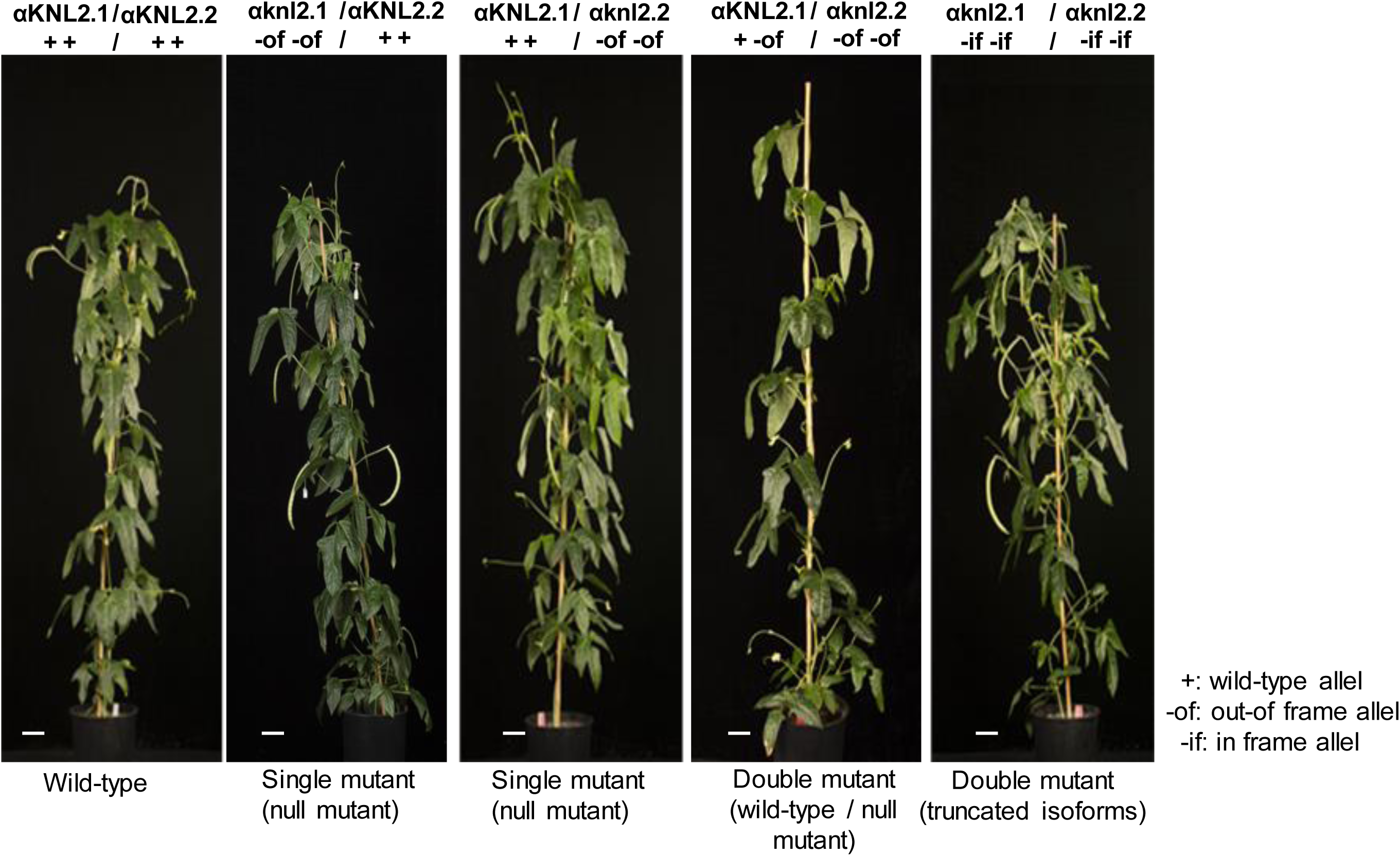
Morphology of various cowpea αknl2 mutants and wild-type. The scale bars correspond to 10 cm.

**Figure S8.**
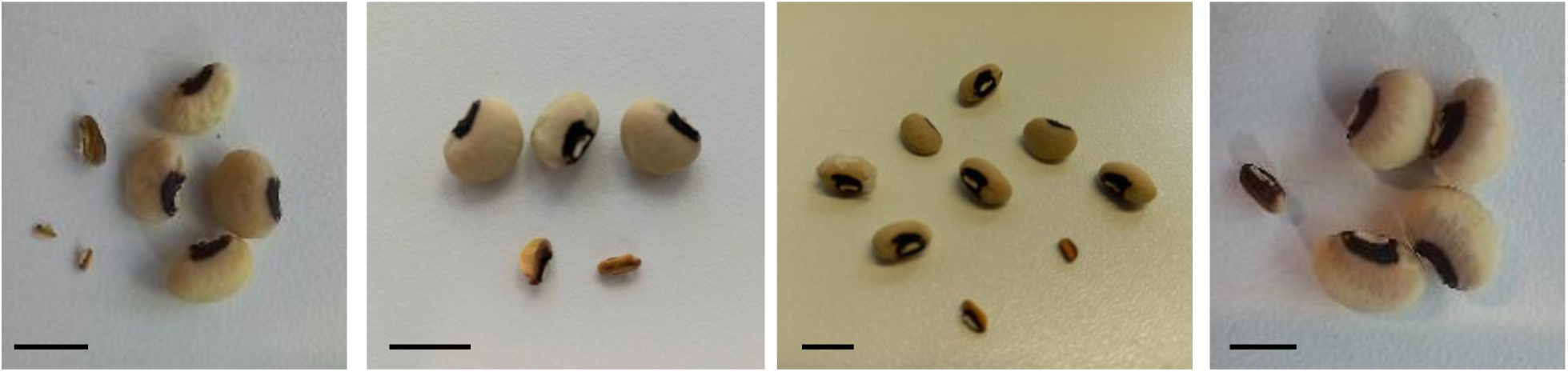
Assay of F3 seeds produced as a result of selfing of plants having knl2.1-of KNL2.1 /knl2.2-of knl2.2-of genotype. Each picture shows the seed set per pod for independent mutant plants; seeds with large-sized “normal” and small-sized “aborted” phenotypes are shown. The scale bars correspond to 5 mm.

**Figure S9.**
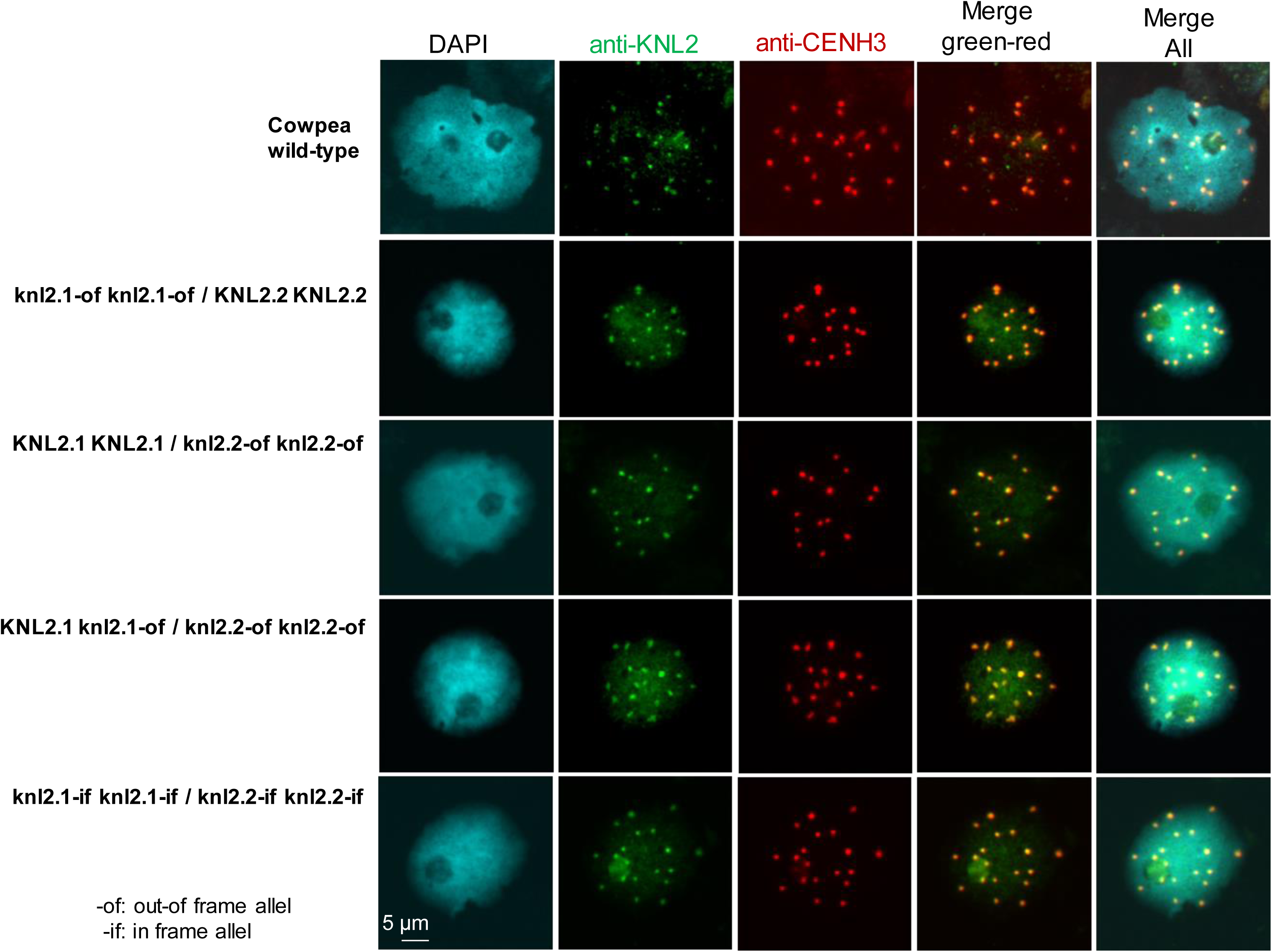
Double immunostaining of mitotic root meristems with anti-αKNL2 and anti-CENH3.1 antibodies. Scale bar 5 µm.

**Figure S10.**
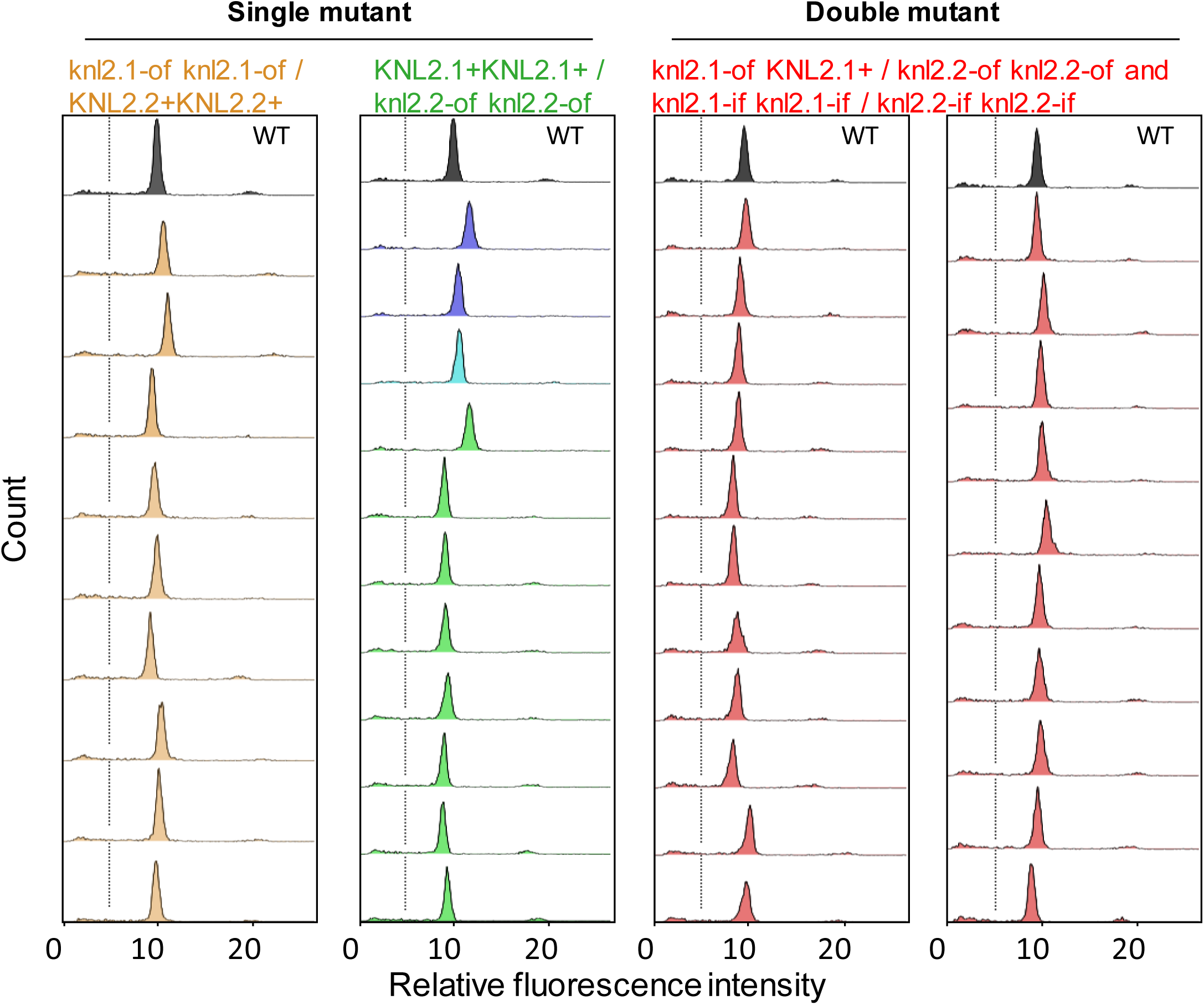
Flow cytometry analysis of F1 seedlings generated as a result of the cross of various αknl2 mutants with wild-type cowpea. The representative curves are presented. Wild-type (WT), black curves; Independent F1 seedlings of the respective mutant, coloured curves.

## Notes

### Competing Interest Statement

The authors have declared no competing interest.

## References

Ahmadli U, Kalidass M, Khaitova LC, Fuchs J, Cuacos M, Demidov D, Zuo S, Pecinkova J, Mascher M, Ingouff M, Heckmann S, Houben A, Riha K, Lermontova I (2023) High temperature increases centromere-mediated genome elimination frequency and enhances haploid induction in Arabidopsis. Plant Commun 4 (3):100507. doi:10.1016/j.xplc.2022.100507

Allipra S, Anirudhan K, Shivanandan S, Raghunathan A, Maruthachalam R (2022) The kinetochore protein NNF1 has a moonlighting role in the vegetative development of Arabidopsis thaliana. Plant J 109 (5):1064–1085. doi:10.1111/tpj.15614

Allu PK, Dawicki-McKenna JM, Van Eeuwen T, Slavin M, Braitbard M, Xu C, Kalisman N, Murakami K, Black BE (2019) Structure of the Human Core Centromeric Nucleosome Complex. Curr Biol 29 (16):2625–2639.e2625. doi:10.1016/j.cub.2019.06.062

Almouzni G, Cedar H (2016) Maintenance of Epigenetic Information. Cold Spring Harb Perspect Biol 8 (5). doi:10.1101/cshperspect.a019372

Altemose N, Logsdon GA, Bzikadze AV, Sidhwani P, Langley SA, Caldas GV, Hoyt SJ, Uralsky L, Ryabov FD, Shew CJ, Sauria MEG, Borchers M, Gershman A, Mikheenko A, Shepelev VA, Dvorkina T, Kunyavskaya O, Vollger MR, Rhie A, McCartney AM, Asri M, Lorig-Roach R, Shafin K, Lucas JK, Aganezov S, Olson D, de Lima LG, Potapova T, Hartley GA, Haukness M, Kerpedjiev P, Gusev F, Tigyi K, Brooks S, Young A, Nurk S, Koren S, Salama SR, Paten B, Rogaev EI, Streets A, Karpen GH, Dernburg AF, Sullivan BA, Straight AF, Wheeler TJ, Gerton JL, Eichler EE, Phillippy AM, Timp W, Dennis MY, O’Neill RJ, Zook JM, Schatz MC, Pevzner PA, Diekhans M, Langley CH, Alexandrov IA, Miga KH (2022) Complete genomic and epigenetic maps of human centromeres. Science 376 (6588):eabl4178. doi:10.1126/science.abl4178

Ariyoshi M, Makino F, Watanabe R, Nakagawa R, Kato T, Namba K, Arimura Y, Fujita R, Kurumizaka H, Okumura Ei, Hara M, Fukagawa T (2021) Cryo-EM structure of the CENP-A nucleosome in complex with phosphorylated CENP-C. The EMBO Journal 40 (5):e105671. 10.15252/embj.2020105671

Bodor DL, Mata JF, Sergeev M, David AF, Salimian KJ, Panchenko T, Cleveland DW, Black BE, Shah JV, Jansen LE (2014) The quantitative architecture of centromeric chromatin. Elife 3:e02137. doi:10.7554/eLife.02137

Boukar O, Belko N, Chamarthi S, Togola A, Batieno J, Owusu E, Haruna M, Diallo S, Umar ML, Olufajo O, Fatokun C (2019) Cowpea (Vigna unguiculata): Genetics, genomics and breeding. Plant Breeding 138 (4):415–424. 10.1111/pbr.12589

Che P, Chang S, Simon MK, Zhang Z, Shaharyar A, Ourada J, O’Neill D, Torres-Mendoza M, Guo Y, Marasigan KM, Vielle-Calzada JP, Ozias-Akins P, Albertsen MC, Jones TJ (2021) Developing a rapid and highly efficient cowpea regeneration, transformation and genome editing system using embryonic axis explants. Plant J 106 (3):817–830. doi:10.1111/tpj.15202

Doyle JJ, Doyle JL (1990) Isolation of DNA from fresh tissue. Focus 12:13–15. 10.2307/2419362

Dunleavy EM, Pidoux AL, Monet M, Bonilla C, Richardson W, Hamilton GL, Ekwall K, McLaughlin PJ, Allshire RC (2007) A NASP (N1/N2)-related protein, Sim3, binds CENP-A and is required for its deposition at fission yeast centromeres. Mol Cell 28 (6):1029–1044. doi:10.1016/j.molcel.2007.10.010

Dunleavy EM, Roche D, Tagami H, Lacoste N, Ray-Gallet D, Nakamura Y, Daigo Y, Nakatani Y, Almouzni-Pettinotti G (2009) HJURP is a cell-cycle-dependent maintenance and deposition factor of CENP-A at centromeres. Cell 137 (3):485–497. doi:10.1016/j.cell.2009.02.040

Engler C, Youles M, Gruetzner R, Ehnert TM, Werner S, Jones JD, Patron NJ, Marillonnet S (2014) A golden gate modular cloning toolbox for plants. Acs Synth Biol 3 (11):839–843. doi:10.1021/sb4001504

Foltz DR, Jansen LE, Bailey AO, Yates JR, 3rd, Bassett EA, Wood S, Black BE, Cleveland DW (2009) Centromere-specific assembly of CENP-a nucleosomes is mediated by HJURP. Cell 137 (3):472–484. doi:10.1016/j.cell.2009.02.039

Fujita Y, Hayashi T, Kiyomitsu T, Toyoda Y, Kokubu A, Obuse C, Yanagida M(2007) Priming of centromere for CENP-A recruitment by human hMis18alpha, hMis18beta, and M18BP1. Dev Cell 12 (1):17–30. doi:S1534-5807(06)00507-7 [pii] 10.1016/j.devcel.2006.11.002

Goodstein DM, Shu S, Howson R, Neupane R, Hayes RD, Fazo J, Mitros T, Dirks W, Hellsten U, Putnam N, Rokhsar DS (2012) Phytozome: a comparative platform for green plant genomics. Nucleic Acids Res 40 (Database issue):D1178–1186. doi:10.1093/nar/gkr944

Grutzner R, Marillonnet S (2020) Generation of MoClo Standard Parts Using Golden Gate Cloning. Methods Mol Biol 2205:107–123. doi:10.1007/978-1-0716-0908-8_7

Hayashi T, Fujita Y, Iwasaki O, Adachi Y, Takahashi K, Yanagida M(2004) Mis16 and Mis18 are required for CENP-A loading and histone deacetylation at centromeres. Cell 118 (6):715–729. doi:10.1016/j.cell.2004.09.002

Hu H, Liu Y, Wang M, Fang J, Huang H, Yang N, Li Y, Wang J, Yao X, Shi Y, Li G, Xu RM (2011) Structure of a CENP-A-histone H4 heterodimer in complex with chaperone HJURP. Genes Dev 25 (9):901–906. doi:10.1101/gad.2045111

Huang H, Strømme CB, Saredi G, Hödl M, Strandsby A, González-Aguilera C, Chen S, Groth A, Patel DJ (2015) A unique binding mode enables MCM2 to chaperone histones H3-H4 at replication forks. Nat Struct Mol Biol 22 (8):618–626. doi:10.1038/nsmb.3055

Ishii T, Juranić M, Maheshwari S, Bustamante FdO, Vogt M, Salinas-Gamboa R, Dreissig S, Gursanscky N, How T, Demidov D, Fuchs J, Schubert V, Spriggs A, Vielle-Calzada J-P, Comai L, Koltunow AMG, Houben A (2020) Unequal contribution of two paralogous CENH3 variants in cowpea centromere function. Communications Biology 3 (1):775. doi:10.1038/s42003-020-01507-x

Jansen LE, Black BE, Foltz DR, Cleveland DW (2007) Propagation of centromeric chromatin requires exit from mitosis. J Cell Biol 176 (6):795–805. doi:10.1083/jcb.200701066

Jefferson RA, Kavanagh TA, Bevan MW (1987) GUS fusions: beta-glucuronidase as a sensitive and versatile gene fusion marker in higher plants. EMBO J 6 (13):3901–3907. doi:10.1002/j.1460-2075.1987.tb02730.x

Jinek M, Chylinski K, Fonfara I, Hauer M, Doudna JA, Charpentier E (2012) A programmable dual-RNA-guided DNA endonuclease in adaptive bacterial immunity. Science 337 (6096):816–821. doi:10.1126/science.1225829

Kozgunova E, Nishina M, Goshima G (2019) Kinetochore protein depletion underlies cyto kinesis failure and somatic polyploidization in the moss Physcomitrella patens. Elife 8. doi:10.7554/eLife.43652

Kozgunova K (2025) Recent advances in plant kinetochore research. Front Cell Dev Biol 12:1510019. doi:doi: 10.3389/fcell.2024.1510019

Lara-Gonzalez P, Pines J, Desai A(2021) Spindle assembly checkpoint activation and silencing at kinetochores. Semin Cell Dev Biol 117:86–98. doi:10.1016/j.semcdb.2021.06.009

Le Goff S, Keceli BN, Jerabkova H, Heckmann S, Rutten T, Cotterell S, Schubert V, Roitinger E, Mechtler K, Franklin FCH, Tatout C, Houben A, Geelen D, Probst AV, Lermontova I (2020) The H3 histone chaperone NASP(SIM3) escorts CenH3 in Arabidopsis. Plant J 101 (1):71–86. doi:10.1111/tpj.14518

Lei Y, Lu L, Liu HY, Li S, Xing F, Chen LL (2014) CRISPR-P: a web tool for synthetic single-guide RNA design of CRISPR-system in plants. Mol Plant 7 (9):1494–1496. doi:10.1093/mp/ssu044

Leite Dias S, Rizzo P, D’Auria JC, Kochevenko A(2025) Efficient Agrobacterium-Mediated Methods for Transient and Stable Transformation in Common and Tartary Buckwheat. Int J Mol Sci 26 (9). doi:10.3390/ijms26094425

Lermontova I, Kuhlmann M, Friedel S, Rutten T, Heckmann S, Sandmann M, Demidov D, Schubert V, Schubert I (2013) Arabidopsis kinetochore null2 is an upstream component for centromeric histone H3 variant cenH3 deposition at centromeres. Plant Cell 25 (9):3389–3404. doi:10.1105/tpc.113.114736

Lermontova I, Schubert V, Fuchs J, Klatte S, Macas J, Schubert I (2006) Loading of Arabidopsis Centromeric Histone CENH3 Occurs Mainly during G2 and Requires the Presence of the Histone Fold Domain. Plant Cell 18 (10):2443–2451

Li Y, Lv M, Gong H, Xu F, Lin Z, Bai Y, Lizhu E, Song W, Lai J, Zhao H (2025) The cenH3 assembly factor ZmKNL2 boosts haploid induction in maize. Plant Biotechnol J 23 (10):4665–4667. doi:10.1111/pbi.70203

Maddox PS, Hyndman F, Monen J, Oegema K, Desai A(2007) Functional genomics identifies a Myb domain-containing protein family required for assembly of CENP-A chromatin. J Cell Biol 176 (6):757–763. doi:10.1083/jcb.200701065

Medina-Pritchard B, Lazou V, Zou J, Byron O, Abad MA, Rappsilber J, Heun P, Jeyaprakash AA(2020) Structural basis for centromere maintenance by Drosophila CENP-A chaperone CAL1. Embo j 39 (7):e103234. doi:10.15252/embj.2019103234

Mellone BG, Fachinetti D (2021) Diverse mechanisms of centromere specification. Curr Biol 31 (22):R1491–R1504. doi:10.1016/j.cub.2021.09.083

Murashige T, Skoog F (1962) A Revised Medium for Rapid Growth and Bio Assays with Tobacco Tissue Cultures. Physiologia Plantarum 15 (3):473–497. doi:DOI 10.1111/j.1399-3054.1962.tb08052.x

Naish M, Alonge M, Wlodzimierz P, Tock AJ, Abramson BW, Schmucker A, Mandakova T, Jamge B, Lambing C, Kuo P, Yelina N, Hartwick N, Colt K, Smith LM, Ton J, Kakutani T, Martienssen RA, Schneeberger K, Lysak MA, Berger F, Bousios A, Michael TP, Schatz MC, Henderson IR (2021) The genetic and epigenetic landscape of the Arabidopsis centromeres. Science 374 (6569):eabi7489. doi:10.1126/science.abi7489

Naish M, Henderson IR (2024) The structure, function, and evolution of plant centromeres. Genome Res 34 (2):161–178. doi:10.1101/gr.278409.123

Pan D, Walstein K, Take A, Bier D, Kaiser N, Musacchio A(2019) Mechanism of centromere recruitment of the CENP-A chaperone HJURP and its implications for centromere licensing. Nature Communications 10 (1):4046. doi:10.1038/s41467-019-12019-6

Parashara P, Medina-Pritchard B, Abad MA, Sotelo-Parrilla P, Thamkachy R, Grundei D, Zou J, Spanos C, Kumar CN, Basquin C, Das V, Yan Z, Al-Murtadha AA, Kelly DA, McHugh T, Imhof A, Rappsilber J, Jeyaprakash AA(2024) PLK1-mediated phosphorylation cascade activates Mis18 complex to ensure centromere inheritance. Science 385 (6713):1098–1104. doi:10.1126/science.ado8270

Pidoux AL, Choi ES, Abbott JK, Liu X, Kagansky A, Castillo AG, Hamilton GL, Richardson W, Rappsilber J, He X, Allshire RC (2009) Fission yeast Scm3: A CENP-A receptor required for integrity of subkinetochore chromatin. Mol Cell 33 (3):299–311. doi:10.1016/j.molcel.2009.01.019

Ranch JP, Liebergesel lM, Garnaat CW, Huffman GA (2010) Auxotrophic Agrobacterium for plant transformation and methods thereof. . WO application WO2010078445A1

Ravi M, Kwong PN, Menorca RMG, Valencia JT, Ramahi JS, Stewart JL, Tran RK, Sundaresan V, Comai L, Chan SWL (2010) The rapidly evolving centromere-specific histone has stringent functional requirements in Arabidopsis thaliana. Genetics 186 (2):461–471. doi:DOI 10.1534/genetics.110.120337

Sato H, Shibata F, Murata M(2005) Characterization of a Mis12 homologue in Arabidopsis thaliana. Chromosome Research 13 (8):827–834

Shelby RD, Monier K, Sullivan KF (2000) Chromatin assembly at kinetochores is uncoupled from DNA replication. J Cell Biol 151 (5):1113–1118. doi:10.1083/jcb.151.5.1113

Silva JdA, Barros JRA, Silva EGF, Rocha MdM, Angelotti F (2024) Cowpea: Prospecting for Heat-Tolerant Genotypes. Agronomy 14 (9):1969

Somasundaram S, Yasar S, Fuchs J, Cuacos M, Claassen J, Weiss O, Kochevenko A, Lamb JC, Li T, Capdeville N, Puchta H, Houben A(2026) Targeted CENH3 protein depletion in egg cells enables highly efficient haploid induction. Plant Commun:101837. doi:10.1016/j.xplc.2026.101837

Spiller F, Medina-Pritchard B, Abad MA, Wear MA, Molina O, Earnshaw WC, Jeyaprakash AA(2017) Molecular basis for Cdk1-regulated timing of Mis18 complex assembly and CENP-A deposition. EMBO Rep 18 (6):894–905. doi:10.15252/embr.201643564

Stankovic A, Guo LY, Mata JF, Bodor DL, Cao XJ, Bailey AO, Shabanowitz J, Hunt DF, Garcia BA, Black BE, Jansen LET (2017) A Dual Inhibitory Mechanism Sufficient to Maintain Cell-Cycle-Restricted CENP-A Assembly. Mol Cell 65 (2):231–246. doi:10.1016/j.molcel.2016.11.021

Takeuchi H, Nagahara S, Higashiyama T, Berger F (2024) The chaperone NASP contributes to de novo deposition of the centromeric histone variant CENH3 in Arabidopsis early embryogenesis. Plant Cell Physiol 65 (7):1135–1148. doi:10.1093/pcp/pcae030

Tamura K, Stecher G, Kumar S (2021) MEGA11: Molecular Evolutionary Genetics Analysis Version 11. Mol Biol Evol 38 (7):3022–3027. doi:10.1093/molbev/msab120

Tello-Ruiz MK, Jaiswal P, Ware D (2022) Gramene: A Resource for Comparative Analysis of Plants Genomes and Pathways. Methods Mol Biol 2443:101–131. doi:10.1007/978-1-0716-2067-0_5

Weber E, Engler C, Gruetzner R, Werner S, Marillonnet S (2011) A modular cloning s ystem for standardized assembly of multigene constructs. PLoS One 6 (2):e16765. doi:10.1371/journal.pone.0016765

Williams JS, Hayashi T, Yanagida M, Russell P (2009) Fission yeast Scm3 mediates stable assembly of Cnp1/CENP-A into centromeric chromatin. Mol Cell 33 (3):287–298. doi:10.1016/j.molcel.2009.01.017

Xie Y, Wang M, Mo B, Liang C (2024) Plant kinetochore complex: composition, function, and regulation. Front Plant Sci 15:1467236. doi:10.3389/fpls.2024.1467236

Yadala R, Camara AS, Yalagapati SP, Vaculikova J, Kralova B, Jaroschinsky P, Meitzel T, Ariyoshi M, Fukagawa T, Potesil D, Rutten T, Palecek JJ, Bui TT, Demidov D, Lermontova I (2026) Structural basis of betaKNL2 centromeric targeting mechanism and its role in plant-specific kinetochore assembly. Nucleic Acids Res 54 (12). doi:10.1093/nar/gkag605

Yalagapati SP, Ahmadli U, Sinha A, Kalidass M, Dabravolski S, Zuo S, Yadala R, Rutten T, Talbert P, Berr A, Lermontova I (2025) Centromeric localization of alphaKNL2 and CENP-C proteins in plants depends on their centromere-targeting domain and DNA-binding regions. Nucleic Acids Res 53 (4). doi:10.1093/nar/gkae1242

Yatskevich S, Yang J, Bellini D, Zhang Z, Barford D (2024) Structure of the human outer kinetochore KMN network complex. Nat Struct Mol Biol 31 (6):874–883. doi:10.1038/s41594-024-01249-y

Zasadzińska E, Barnhart-Dailey MC, Kuich PH, Foltz DR (2013) Dimerization of the CENP-A assembly factor HJURP is required for centromeric nucleosome deposition. Embo j 32 (15):2113–2124. doi:10.1038/emboj.2013.142

Zasadzińska E, Huang J, Bailey AO, Guo LY, Lee NS, Srivastava S, Wong KA, French BT, Black BE, Foltz DR (2018) Inheritance of CENP-A Nucleosomes during DNA Replication Requires HJURP. Dev Cell 47 (3):348–362.e347. doi:10.1016/j.devcel.2018.09.003

Zhang Z, Bower F, Komaki S, Pettko-Szandtner A, Skuba AO, Tromer EC, Magyar Z, Heese M, Schnittger A(2025) Functional characterization of the Csm1-like protein TITAN 9 in Arabidopsis thaliana. iScience 28 (9):113251. doi:10.1016/j.isci.2025.113251

Zuo S, Yadala R, Yang F, Talbert P, Fuchs J, Schubert V, Ahmadli U, Rutten T, Pecinka A, Lysak MA, Lermontova I (2022) Recurrent plant-specific duplications of KNL2 and Its conserved function as a kinetochore assembly factor. Mol Biol Evol 39 (6). doi:10.1093/molbev/msac123

